# FGF8 induces chemokinesis and regulates condensation of mouse nephron progenitor cells

**DOI:** 10.1101/2022.02.07.478973

**Authors:** Abhishek Sharma, Marco Meer, Arvydas Dapkunas, Anneliis Ihermann-Hella, Satu Kuure, Seppo Vainio, Dagmar Iber, Florence Naillat

## Abstract

Kidneys develop via iterative branching of the ureteric epithelial tree and subsequent nephrogenesis at the branch points. Nephrons form in the cap mesenchyme as the metanephric mesenchyme (MM) condenses around the epithelial ureteric buds (UBs). Previous work demonstrated that FGF8 is important for the survival of nephron progenitor cells (NPCs), and early deletion of *Fgf8* leads to the cessation of nephron formation, which results in post-natal lethality. We now reveal a novel function of FGF8. By combining transgenic mouse models, quantitative imaging assays, and data-driven computational modelling, we show that FGF8 has a strong chemokinetic effect and that this chemokinetic effect is important for the condensation of NPCs to the UB. The computational model shows that the motility must be lower close to the UB to achieve NPC attachment. We conclude that the FGF8 signalling pathway is crucial for the coordination of NPCs behaviour to the UB. Chemokinetic effects have been described also for other FGFs and may be relevant more generally for the formation of mesenchymal condensations.

## Introduction

Mesenchymal condensation is an essential step in kidney development for the early formation of nephrons. This mechanism consists of reciprocal interactive signalling between mesenchymal cells and their surrounding, the epithelial and stromal cells [1–3]. In addition to reciprocal signalling, intercellular interactions, cellular morphogenesis, i.e. apoptosis or adhesion, and cell migration play an essential role during the establishment of mesenchymal condensation [4–6]. Cell migration can be influenced by chemical, thermal, galvanic, electrical, gravitational, or mechanical stimuli, or combinations of these phenomena. A stimulus can cause a tactic response, in which cell movement is directed to the location of the stimulus, or a kinetic response, i.e., random locomotion, in which the magnitude of the response depends on the intensity of the stimulus [7]. Particularly in the presence of chemical gradients, cells can show strong chemotactic or chemokinetic responses. In mice, around embryonic days 11-11.5 (E11-E11.5), mesenchymal condensation in the nephrogenic niche of the developing kidney results in the formation of the cap mesenchyme (CM) [2, 8]. At the same time, nephron progenitor cells (NPCs) from the CM migrate in a stochastic fashion between the top (or tip region) of the epithelial ureteric bud (UB) and the bottom (or trunk region) (Figure 1) [9–11]. NPC fate is region-specific and requires reciprocal signals between the UB and the surrounding mesenchymal and stromal cells (Figure 1) [1, 2, 11–16]. NPCs that are located in the tip region of the UB maintain their progenitor state and are thus called true nephron progenitor cells (tNPCs) (Figure 1) [10, 14, 17]. NPCs that migrate downwards to the trunk region are further primed by factors from the UB, becoming committed NPCs (cNPCs) [2, 12, 14, 15]. A fraction of the cNPCs in the trunk region starts to form a pretubular aggregate (PTA), initiating nephron formation [2, 9, 11, 12, 18]. The regulation of NPC fates, migration, and priming have been studied intensely [2, 3, 19], but the cellular mechanism underlying the condensation of NPCs to the UB has not yet been elucidated. Various signaling factors, receptors, and extracellular matrix molecules have been suggested to play a role in NPC condensation (Figure 1), but its key regulators remain elusive [9, 20–22].

**Figure 1:**
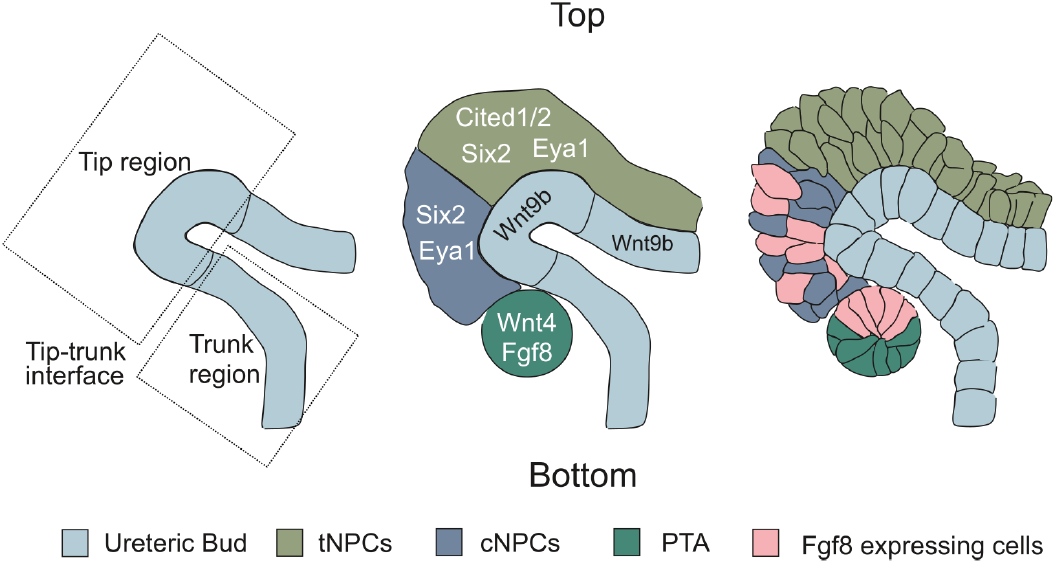
Cartoon depicting the UB and the CM nephron progenitor population during early kidney development. tNPCs express *Cited1/2*, *Six2* and *Eya1*, while cNPCs lose *Cited1/2* expression upon priming by Wnt9b from the UB. A subset of cNPCs continues to form a PTA at the tip-trunk interface. Schematic based on [10, 17, 23].

Fibroblast growth factors (FGFs) are a family of signaling proteins that govern different aspects of kidney development, including UB branching and maintenance of NPCs [24]. Deletion or mutations in either FGFs or their receptors (FGFRs) can lead to either kidney agenesis or disorders [24]. FGF8 is expressed in the mesenchyme and is required for both the regulation of downstream genes involved in PTA formation and cell survival [12, 25–27]. Deletion of *Fgf8* from kidney primordia leads to a lack of mature nephrons, and eventually to lethality within 24hrs of birth [25]. The failure of nephron maturation has been attributed to the lack of expression of *Wnt4* and *Lim1 (Lhx1)*, both of which are crucial for MET [25]. Culturing isolated MM from kidneys lacking *Fgf8* along with a WNT source (embryonic spinal cord), however, failed to initiate nephrogenesis [25, 26], supporting the notion that WNT4 and FGF8 work independently. Further, when ectopic FGF8 was added in combination with a WNT source, MMs lacking *Fgf8* expression formed PTAs [25]. These results indicate that FGF8 enhances WNT4 expression in PTAs but also suggest that FGF8 and WNT4 work independently. As little is known about the specific role of FGF8 during cap mesenchyme formation, we further characterise its specific role to demonstrate that the FGF8 signal coordinates NPC migration during mesenchymal condensation.

## Results

### Without the expression of *Fgf8*, cap mesenchyme formation and attachment of CM cells to the UB are impaired

When the kidney develops from the posterior intermediate mesoderm [28], the *Brachyury/T* gene is required for the formation of the posterior mesoderm and axial development [29]. Hence, *Brachyury/T^Cre^*-mediated deletion leads to the deletion of *Fgf8* in both epithelial and mesenchymal compartments of the developing kidney [25, 28, 29]. To examine more closely the involvement of FGF8 in kidney development we stained mutant kidneys (*Fgf* 8^*n/c*^;*T^Cre^*) with SIX2, a known NPC marker [2, 3]. In *Fgf* 8^*n/c*^;*T^Cre^* mutant kidneys, we found that the *SIX*2^+^ cell population was diminished and less condensed than in the controls (Figure 2A,B). The reduced cell number has previously been attributed to increased cell death in *Fgf8* -deficient kidneys [25]. We hypothesized that the absence of FGF8 signaling additionally leads to decreased cell motility and consequent failure of cell condensation and eventually the termination of nephrogenesis. To test this hypothesis, we first investigated whether the failure of *SIX*2^+^ cells to condensate would also occur in an *in vitro* culture assay. We used the Trowell culture method to culture *Fgf* 8^*n/c*^; *T^Cre^* kidneys and littermate control kidneys. After 3 days of culture, corresponding to E16.5, we found that the *SIX*2^+^ cells in mutant kidneys were significantly less condensed as compared to their littermate controls (Figure 2C-D). More specifically, we found that the CM in mutant kidneys was on average twice as thick as in the controls (Figure 2E). In accordance with this observation, the *SIX*2^+^ cells in mutant kidneys were strongly dispersed within the niche (Figure 2F). We therefore wondered whether the number of *SIX*2^+^ cells that are attached to the UB was also affected as a result. Attachment to the UB has previously been suggested to affect the differentiation capacity of mesenchymal cells [15]. We classified *SIX*2^+^ cells to be attached to the UB when their centroid position was closer to the UB tip than a threshold value of 17.4 μm, which corresponds to twice the median position of *SIX*2^+^ cells that we measured in the controls (Figure 2F). Our choice of threshold value agrees well with the upper quartile of distances that were previously ascribed to the attached state (15.2μm, [9]). We also found that the percentage of attached cells in our controls (92%, Figure 2G) was in good agreement with what was previously found (85%, cf. Figure 5 source data of [9]). Finally, we found that in our *Fgf8* -deficient mutant kidneys the proportion of *SIX*2^+^ cells that was attached to the UB tip was significantly lower than in the controls (Figure 2G). Thus, as a next step, we wanted to investigate the effect of FGF8 on the *SIX*2^+^ NPC population in more detail.

**Figure 2:**
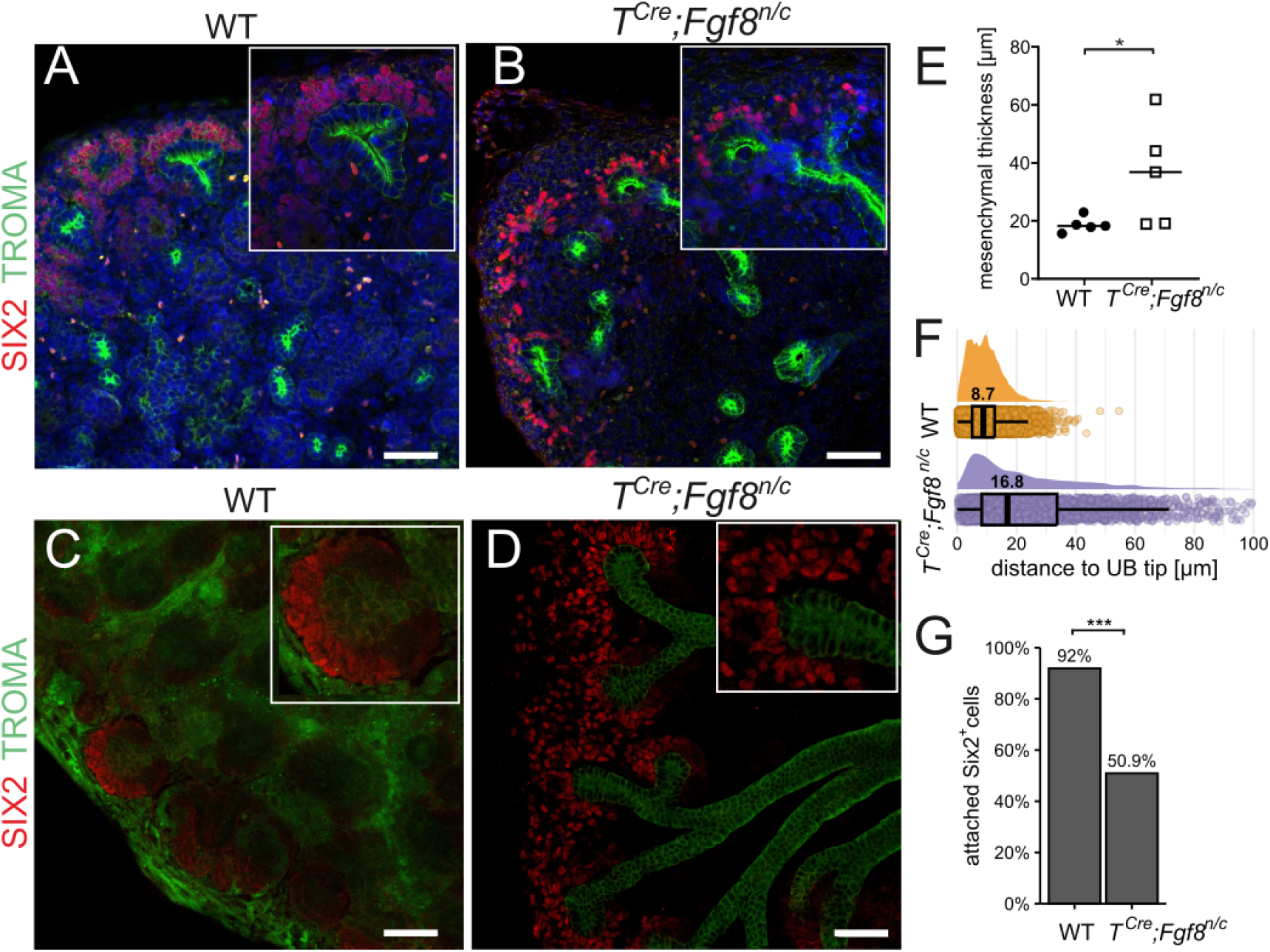
Impaired cap mesenchyme formation. (A-D) Kidneys were stained with SIX2 (NPCs, red), Troma-1 (UB, green) and DNA was counter stained with Hoechst33342 (Blue). Scale bars represent 100 μm. (A) E16.5 littermate control. (B) E16.5 kidneys where *Fgf8* was deleted using *T^Cre^*. (C) Littermate control E11.5 kidneys cultured in Trowell culture system. (D) E11.5 kidneys where *Fgf8* was deleted using *T^Cre^* cultured in Trowell system. (B,D) NPCs did not form a condensed cap mesenchyme, but were scattered around the tip region of the UB as compared to the littermate controls (A,C), respectively. The inserts are higher magnification selections. (E) Cap mesenchymal thickness as the distance of the last layer of *Six*2^+^ cells from the UB tip of cultured E11.5 *Fgf* 8^*n/c*^; *T^Cre^* kidneys. Statistics were performed in the kidneys (n=6) per sample, *p* values were calculated using *t-test*, *p < 0.05. (F) Distributions of distance between centroids of *Six*2^+^ cells and UB tip in E16.5 Trowell cultured *Fgf* 8^*n/c*^; *T^Cre^* kidneys (n=13) and littermate controls (n=15), pooled data. Numbers above the boxplots represent median distances. (G) Percentage of *Six*2^+^ cells attached to the UB. Total number of cells (pooled data): 11425 (controls), 3214 (*Fgf* 8^*n/c*^; *T^Cre^*). *Six*2^+^ cells were classified as attached to the UB when their centroid position was within a distance of 17.4 μm from the UB, corresponding to twice the median position of *Six*2^+^ cells in the controls. Significance level: *** p < 0.001.

### FGF8 induces NPC aggregate formation *in vitro* and is required for tNPC maintenance

The *SIX*2^+^ NPCs dominate the CM population around the UB [30], and when these cells are primed as cNPCs, a subset of these cNPCs forms a PTA [2, 3]. In an exhaustive study of secreted FGF family members which are required for the maintenance of the tNPC state, it has been suggested that FGF8 fails to maintain the true progenitor state of NPCs [31]. However, this study made use of a 2D monolayer culture, which differs from the 3D *in vivo* microenvironment of the developing kidney. Ihermann-Hella *et al.* and Dapkunas *et al.* have since developed a 3D culture system for NPCs [32, 33]. Much as in the 2D cultures, MM cells form aggregates when cultured with FGF2 in the 3D cultures [32, 34]. To test whether NPCs would also condensate in response to FGF8, we cultured the dissociated NPCs from E11.5 kidneys in a 3D matrix along with FGF2 (positive control), or FGF8, or anti-FGF8 antibody to block any FGF8 secreted by MM cells. The chosen monoclonal anti-FGF8 antibody binds selectively to FGF8 in *in vitro* experiments (Supplementary Figure 1). From dissected E11.5 kidneys, MM was separated from UB, dissociated into single cell suspension, seeded in matrigel and cultured for 24 hours.

In response to both FGF2 and FGF8, the NPCs formed condensates and retained *Six2* expression (Figure 3B,C), while both the control and NPCs treated with anti-FGF8 antibody lost SIX2 expression (Figure 3A,D, Supplementary Movie 1). We also found that the loss of SIX2 expression as a result of the absence of FGF8 (Figure 3A,D) was not completely due to cell death as more live cells were observed (Figure 3E). This suggests that FGF8 plays an additional role in NPC commitment along with its role in forming NPC condensates. To examine the differentiation stage of NPCs between tNPCs and cNPCs, a qPCR analysis of NPCs in the presence or absence of FGF8 was carried out. tNPC markers such as *Cited2*, *Six2* and *Eya1* were maintained by ectopic FGF8 (Figure 3F). To confirm that the results were not influenced by signals from the UB, the same experiment was carried out with fully dissociated MMs lacking UB. Similar results were obtained with retained tNPC markers when treated with ectopic FGF8 (Fig 3G). It has been reported that cross-talk between WNT and FGF signalling pathways is linked by modulation of phosphorylation of *GSK*3*β* [35], but we did not observe this in our setting (Supplementary Figure 2). Finally, in wild type kidneys that were cultured in the presence of ectopic FGF8 (Trowell culture method), we could observe an expansion of the *SIX*2^+^ population (Figure 3H). In conclusion, our 3D culture matrix experiments suggest that FGF8 is required for both NPC condensation and tNPC maintenance.

**Figure 3:**
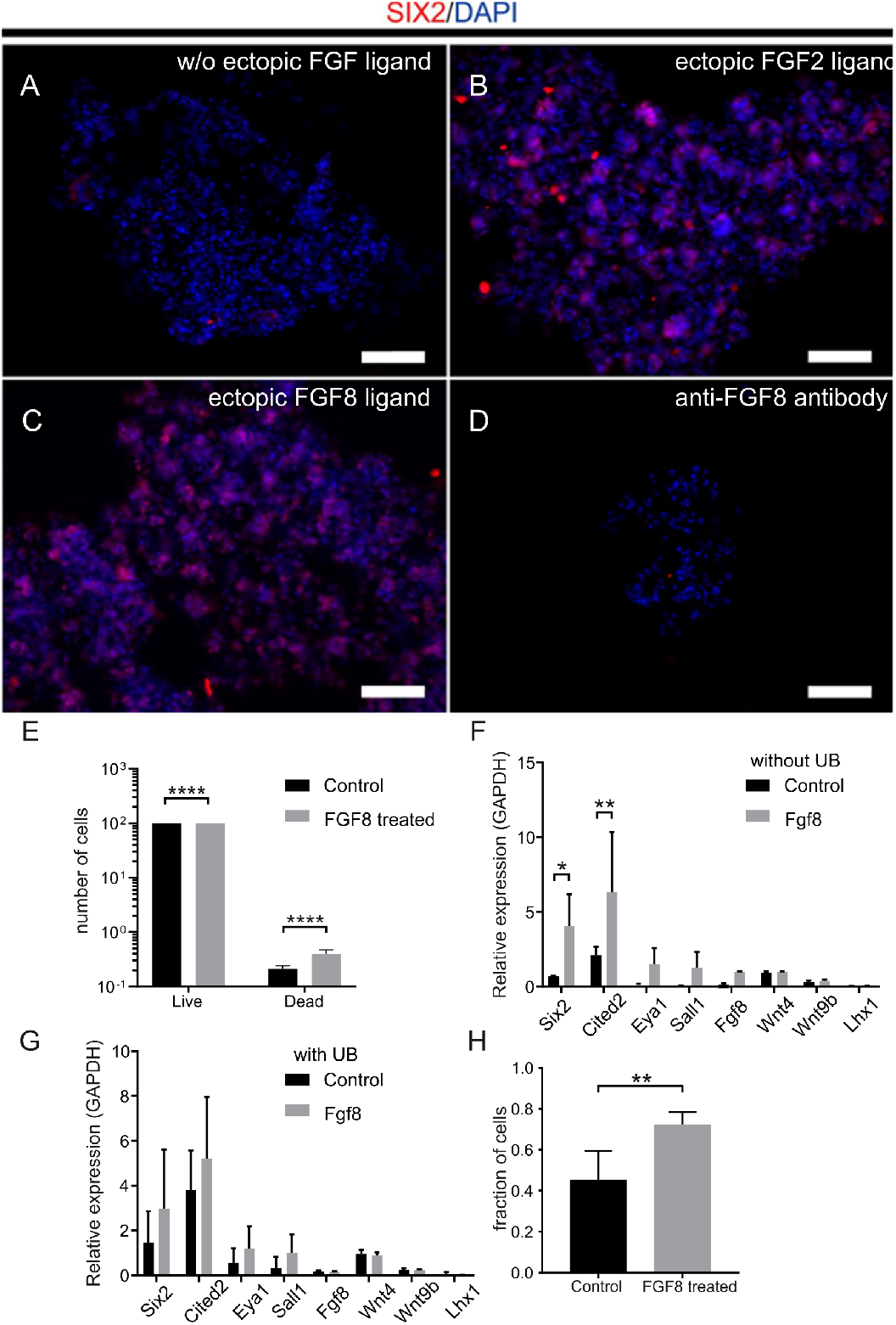
Gene expression analysis in NPCs in the presence or absence of FGF ligand. (A-D) Antibody staining of SIX2 expression with DAPI counterstaining. (A) NPCs lose the expression of SIX2 when the ectopic FGF ligand is not provided. (B,C) NPCs aggregate and retain SIX2 expression in the presence of FGF2 (B) or FGF8 (C). (D) Loss of SIX2 expression and cell death in the presence of anti-FGF8 antibody. (A-D)Scale bar represents 100 μm. (E) Loss of SIX2 expression and cell death in NPCs treated with anti-FGF8b antibody analysed by flow cytometry. Live/dead analysis was performed using flow cytometry (7-AAD staining), statistics (n=3) were performed based on two-way ANOVA with significance level ****p < 0.0001. (F,G) qPCR of tNPC gene expression in FGF8-treated nephrospheres (F) and in the presence of the UB (G). Statistics (n=3) were performed based on t-test, where *p < 0.01 and **p < 0.001; if not specified, the test results were not significant. (H) Percentage of *SIX*2^+^ cells in response to ectopic Fgf8. Quantitative flow cytometry of dissociated E11.5 kidneys. Samples were collected using FACSCalibur and analyzed using FlowJo, statistics (n=10) were performed based on t-test, **p < 0.001.

### FGF8 elicits a chemokinetic response

FGFs have been shown to act as chemoattractants that trigger a chemotactic response [37, 38], i.e., the migration of a cell along the concentration gradient of the chemoattractant. To determine how FGF8 affects NPC motility, we placed an FGF8 soaked bead in a 3D matrigel matrix containing MM cells from E11.5 kidneys and tracked cells (Supplementary Movie 2). Cell tracking revealed that MM cells migrated significantly faster compared to the control bead experiments, but cellular motion was stochastic and lacked directionality (Figure 4; Supplementary Movie 2). This shows that FGF8 has mainly a chemokinetic effect, i.e., an impact on the speed of movement, rather than a chemotactic effect. Here, we note that we did not measure the FGF8 concentration profile and thus cannot exclude that meaningful gradients did not emerge due to rapid dispersion of FGF8 in matrigel. Previous measurements in zebrafish showed that the diffusion coefficient of FGF8 is high in aqueous environments [39]. The measured cell speeds (4.6 *±* 0.5 μm/h; Figure 4 N) are comparable to those measured by others in the UB tip region [9, 40]. Interestingly, in the presence of FGF8 soaked beads, most cells formed large aggregates, resulting in a collective movement resembling swarm behaviour, which is reflected in the bifurcation of the track straightness measurements of FGF8 vs. control (Figure 4 E,F, Supplementary Movie 2). In the control bead experiments, only a few small aggregates were formed, presumably due to low levels of *Fgf8* expression by some MM cells.In summary, we find that FGF8 triggers a chemokinetic response of MM cells from E11.5 kidneys. This observation of undirected cellular motility agrees with previous quantifications where MM cells were found to move semi-stochastic and swarming-like [9].

**Figure 4:**
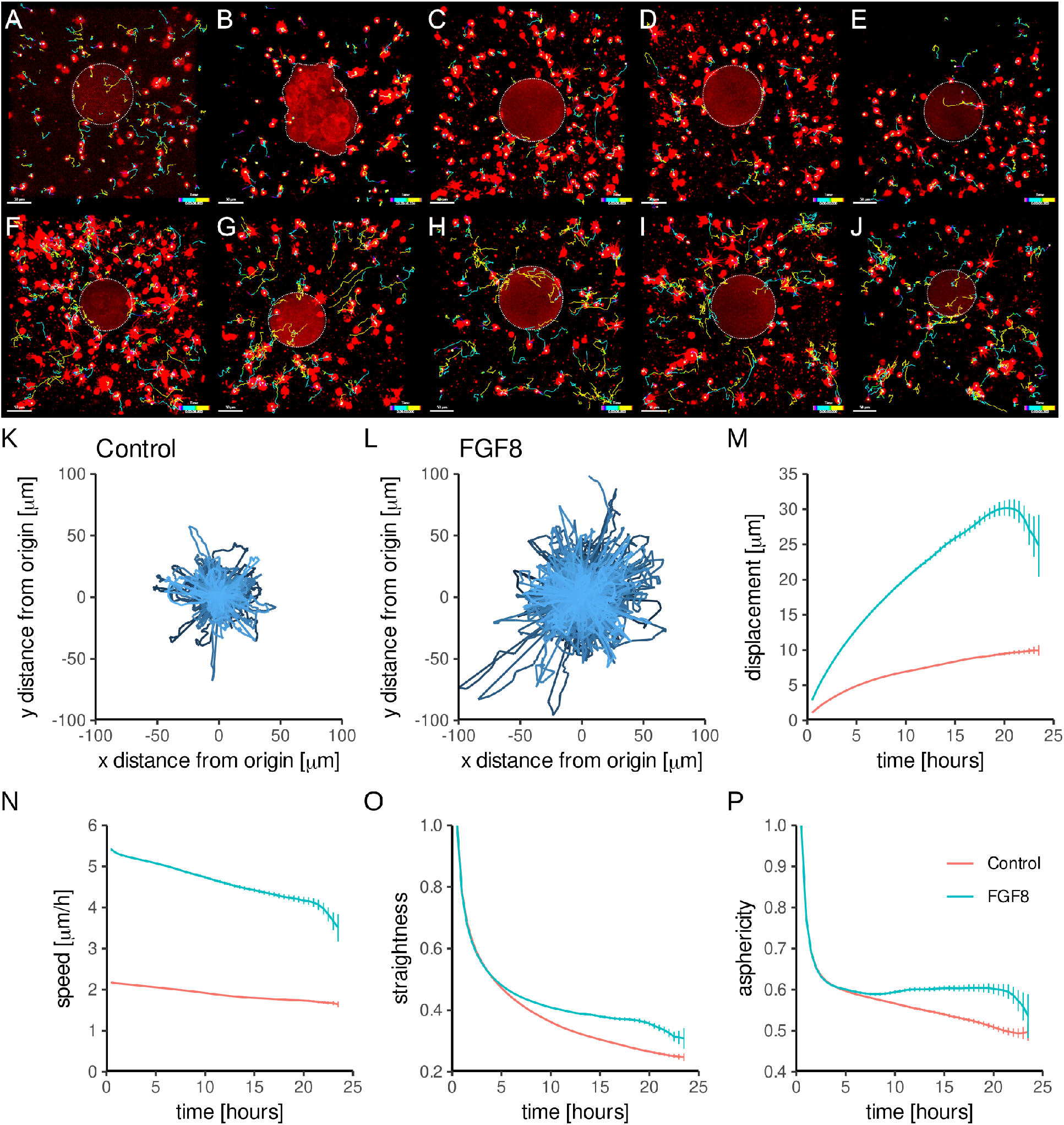
Elevated and undirected NPC motility in response to FGF8. *In vitro* experiments with control beads (A-E) vs. FGF8 soaked beads (F-J), in a culture of E11.5 MM cells, 5 samples each. Full cell tracks are shown after 24-hour time lapse (FGF8: 534 tracks *in toto*, control: 538 tracks *in toto*). Cells were tracked for 24hrs. Beads are highlighted by dotted white lines. Scale bar represents 50 μm. Tracks from all samples of each of the two cohorts were pooled (FGF8: 534 tracks, control: 538 tracks). Star plots of control (K) and (L) FGF8 experiments were normalized such that the start point of each track was at the origin. (M,N) Average displacement (Euclidean distance between track start and endpoints) and average speed over time. (O,P) Track straightness: Displacement divided by contour length. Asphericity: The less spherically the steps of a track are distributed, the straighter the track (cf. K,L and [36]).

### A model based on FGF8-induced motility leads to robust condensation of NPCs

We next thought to test the impact of FGF8-induced chemokinesis on NPC condensations at the UB using computational modelling. UB outgrowth starts at around E10.5 when the metanephric mesenchyme is in a diffuse, thickened state, inducing the patterning of the MM [8, 12, 34, 41–43]. At around E11, diffuse weak expression of *Fgf8* by MM cells coincides with the emergence of a well-defined cap mesenchyme [8, 12, 43]. Close to the UB, MM cells are rather immotile [9]. Interestingly, Sonic hedgehog (SHH), a repressor of *Fgf8* expression, is secreted from UB cells during early nephrogenesis [39, 44]. The SHH gradient emanating from the UB likely results in the lower *Fgf8* expression that is observed closer to the UB [45]. A gradient of autocrine FGF8 signaling and thus chemokinesis would be in agreement with the previous observation that cell speeds increase with distance from the UB [9]. In the same study, it was found that MM cells experience a subtle attraction towards the UB, indicating the presence of a chemotactic factor. WNT11 represents a likely candidate, as it is secreted already around E10.5-E11 by UB tip cells and is required for stable NPC-UB attachment [12, 15, 46, 47].

To analyze the interplay of chemical signaling and cell motility during mesenchymal condensation, we asked whether a model consisting of i) FGF8-induced mesenchymal cell motility, ii) WNT11-based chemoattraction and iii) SHH-induced repression of FGF8 close to the UB can explain mesenchymal condensation around the UB (Figure 5A). In the computational model, NPCs are initially randomly positioned in a niche bordering a flat patch of the ureteric epithelium (Figure 5E). The NPCs are assigned velocities that depend on the local FGF8 concentration, while the direction of movement is chosen randomly (Figure 5A,B). Epithelial cells are secreting weak concentrations of both WNT11 and SHH, while scenarios of different levels of *Fgf8* expression by MM cells are explored (Figure 5F,G; Supplementary Figure 3). Mechanically, all cells can adhere to each other when being closer than a threshold distance and detach when moving apart. Simulating this model results in an effective motility gradient with a trap-like region close to the epithelium (Figure 5C,D,F; Supplementary Figure 3). At weak concentrations of FGF8, WNT11 and SHH, only the MM cells closest to the ureteric epithelium show strong displacement towards the ureteric epithelium, but most cells that are further away remain within a few cell diameters of their initial positions (Figure 5C,F; Supplementary Movie 3). At intermediate (2x increased) concentrations of FGF8, significantly more cells aggregate close to the basal surface of the ureteric epithelium (Figure 5C,D,G). At higher FGF8 concentrations (3x increased), cells aggregate everywhere in the niche, which is in line with our bead experiments (Figure 5C, Supplementary Movie 3; Supplementary Movie 2). Lastly, we also observe FGF8 peaks at locations where cells aggregate, resulting in swarm-like motility.

**Figure 5:**
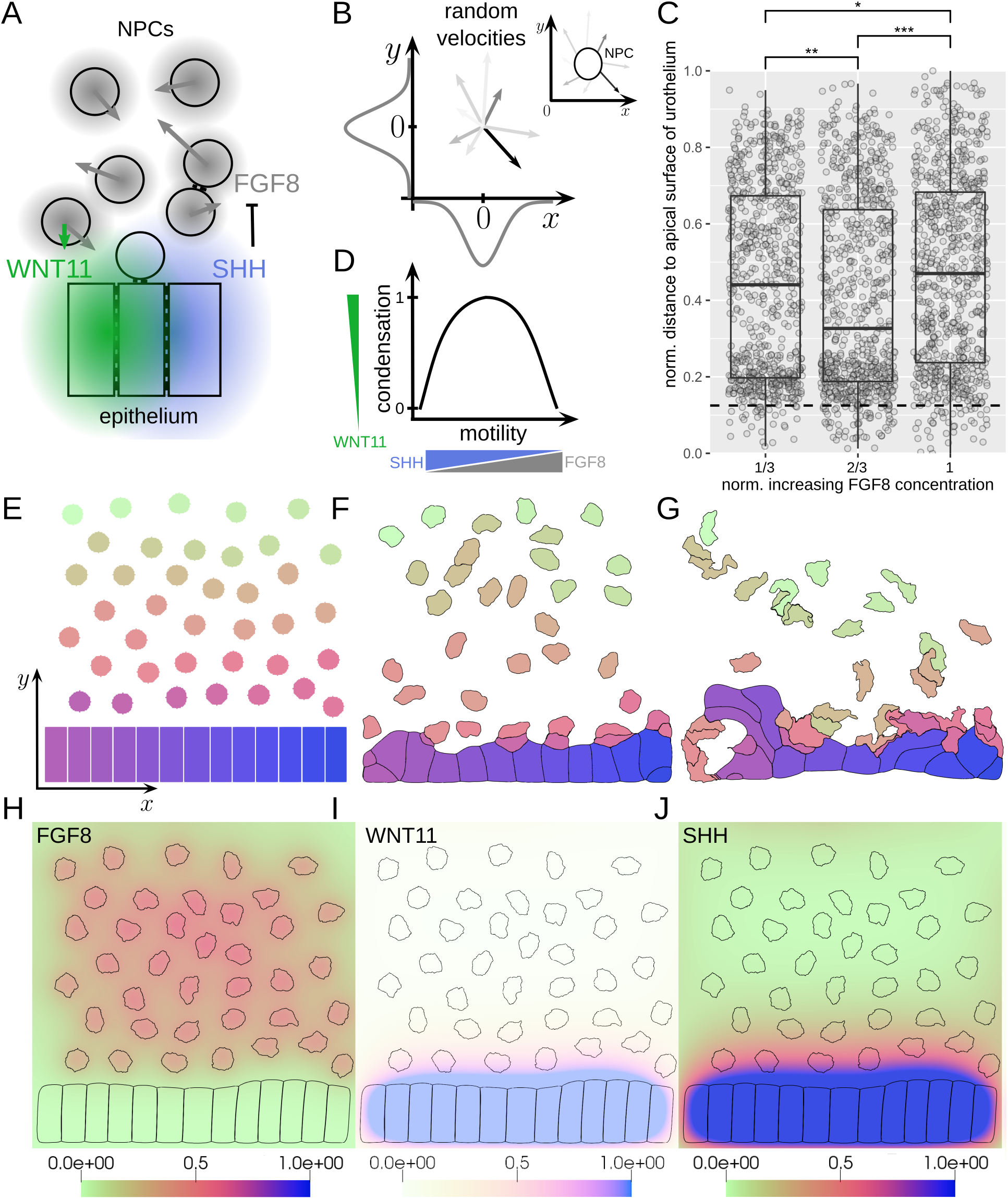
2D simulations of NPC condensation to the ureteric epithelium. (A) Cartoon of the simulation setup. NPCs express FGF8, can adhere to each other and the ureteric epithelium and move with random velocities whose magnitude depends on FGF8 concentration. Ureteric epithelial cells also adhere to each other and secrete SHH that represses *Fgf8* expression, and WNT11, which induces a weak attraction towards the ureteric epithelium (with one order of magnitude lower NPC speeds than with FGF8). NPCs close to the ureteric epithelium are immobilized due to SHH-induced repression of *Fgf8*. (B) Implementation of random cell motility. The components of the 2-dimensional velocity vector for each cell GeometryNode are picked from normal distributions. This implementation better reflects the amoeboid shape-shifting behaviour of the NPCs. Inlet: Example cell with velocities for a few GeometryNodes. In the simulations, each cell possesses several hundred GeometryNodes. Cf. methods section. (C) Distance of NPCs to ureteric epithelium for simulations with an increasing FGF8 concentration. Only intermediate levels of FGF8 lead to more NPCs being trapped in the region close to the basal surface of the ureteric epithelium (dashed line). Data were pooled from n=20 simulations for each group. Significance levels: *p < 0.1, **p < 0.01, ***p < 0.001. (D) Motility decreases with an increasing concentration of SHH and increases with increasing concentrations of FGF8 and WNT11. Cells quickly condensate with increasing FGF8 concentrations. In the simulations, the WNT11 concentrations are one order of magnitude smaller than the FGF8 concentrations and accordingly, cell speeds (Table 4, Supplementary Figure 3). (E) Start configuration with dispersed NPCs over a flat patch of the ureteric epithelium. Colours correspond to cell identity. (F) Example of a final configuration with low average FGF8 concentration and limited NPC displacement. (G) Example of a final configuration with an intermediate FGF8 concentration, with increased NPC displacement and mixing. (H-J) Examples of initial concentration gradients of FGF8, WNT11 and SHH. Colour codes represent normalized concentrations.

### Deletion of *Fgf8* in late nephrons leads to hypomorph kidney phenotype

Deletion of *Fgf8* before gastrulation is lethal as cells lose the ability to migrate away from the primitive streak [25, 48]. Deletion of *Fgf8* in the MM using *Pax*3^*Cre*^ mice leads to the same phenotype, and newborn mice die shortly after birth [26]. We wondered whether FGF8 also plays a role in later stages of kidney development. *In situ* hybridisation of *Fgf8* revealed its expression in upper cells forming PTAs and this expression was still maintained in the top cells of renal vesicles that form the future comma and S-shape bodies in WT kidneys (Figure 6A-E). To investigate its effect during early nephron formation at PTA stage, we used later expression tissue-specific Cre mouse lines (described in Material and Methods section). Cre was under the promoter of *Wnt4* or *Pax8* genes. By utilizing this strategy, we generated mutant *Fgf* 8^*n/c*^ progeny with 50% frequency, while *Fgf* 8^*n/*+^ mice were used as littermate controls. First, to be sure that *Fgf8* did not have any function in the UB, we first deleted *Fgf8* from the UB using *HoxB*7^*Cre*^ mice, and as expected the mutant had no phenotype and the kidneys were similar to those of the wild type mice (Supplementary Figure 4C, Supplementary Figure 5A-D [49]).

**Figure 6:**
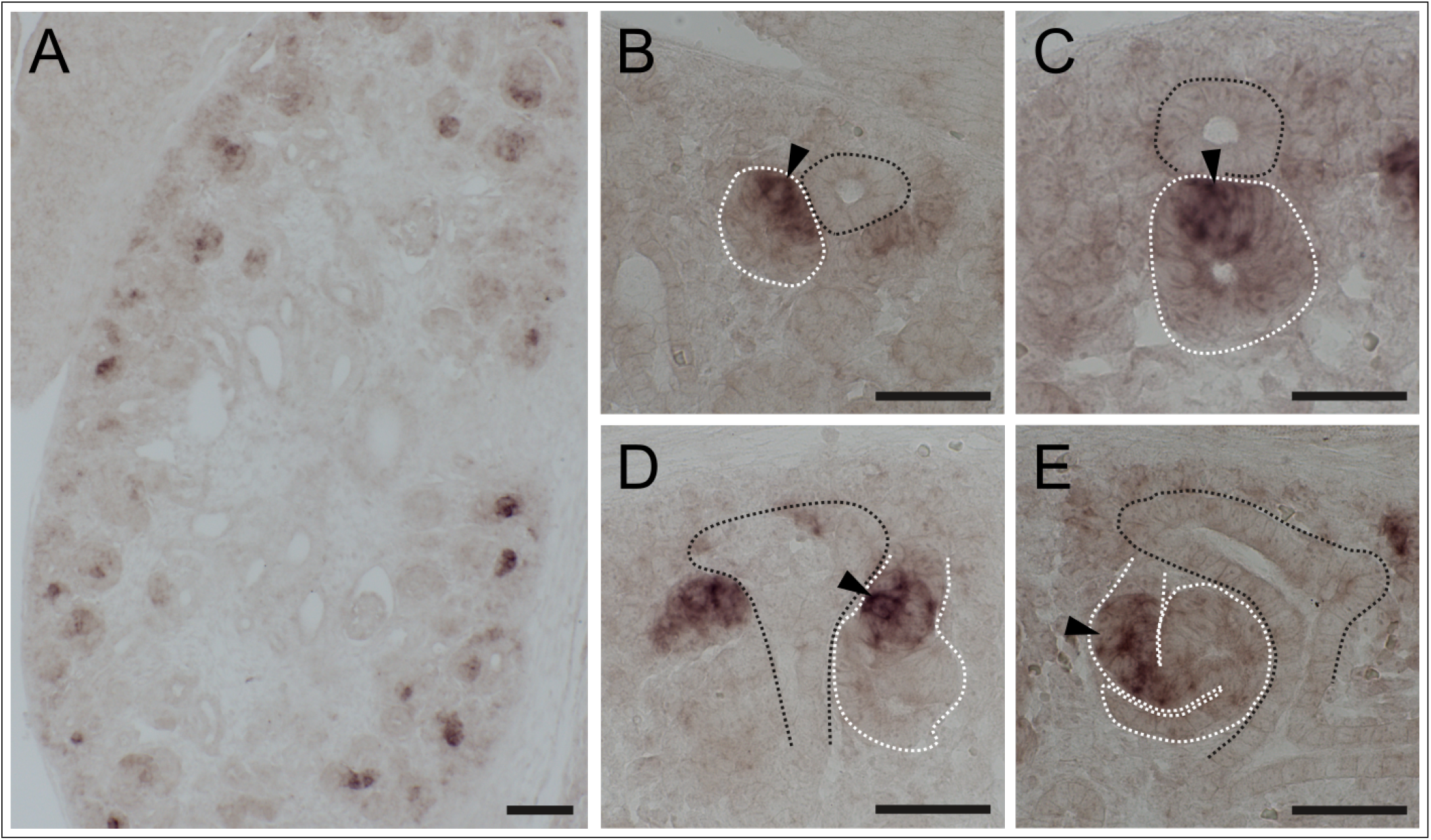
*In situ* hybridization of *Fgf8* in E15.5 embryonic mouse kidney. Taking a closer look at MM derivatives (B-E), the expression of *Fgf8* is localized closer to the UB tip. (A) 10x magnification of E15.5 kidney. (B-E) 43x magnified selections. Dotted black lines indicate the ureteric epithelium, dotted white lines indicate (B) pretubular aggregate, (C) renal vesicle, (D) Comma-shaped body and (E) S-shaped body. Black arrows indicate cells expressing *Fgf8*. Scale bar for (A) 100 μm and (B-E) 50 μm.

Second, we deleted *Fgf8* from MM cells by employing two tissue-specific Cre recombinase mice (under the promoter of *Pax8* and *Wnt4* gene). *Pax*8^*Cre*^ is expressed in both the MM and the UB tip (GUDMAP; Supplementary Figure 4A) [50, 51], and *Wnt*4^*Cre*^ is only expressed in the MM (Supplementary Figure 4B [52]). We found that the *Fgf8* deletion from the MM using either *Pax*8^*Cre*^ or *Wnt*4^*Cre*^ led to smaller kidneys as compared to littermate controls (Figure 7A-C). On closer inspection, we found that in both cases kidneys had fewer mature nephrons as compared to littermate controls. Kidneys of *Fgf8*; (*Pax*8^*Cre*^) revealed an arrest in S-shaped-body structures whereas the *Fgf8* ; (*Wnt*4^*Cre*^) showed comma-shaped body structures (Figure 7D-I, higher magnification). Because the complete loss of FGF8 function during kidney development results in a failure of nephron formation around the S-shaped body stage [25, 26], these results raised the question is there still some *Fgf8* expressed in the *Fgf* 8^*n/c*^;*Pax*8^*Cre*^ and *Fgf* 8^*n/c*^;*Wnt*4^*Cre*^ kidneys? Therefore, we analyzed the *Fgf8* expression in *Fgf* 8^*n/c*^;*Pax*8^*Cre*^ kidneys. Functional *Fgf8* RNA is expressed at lower level than control litter analysed by qPCR suggesting that the remaining *Fgf8* expression cause an hypomorph kidney phenotype. We observed around 40% of *Fgf8* expression at E12.5 in kidneys lacking *Fgf8* (Figure 7J). These data demonstrate that FGF8 is required within the developing kidney to support the further development of the nephrons, and the reduced Fgf8 expression during nephrogenesis induces hypomorph phenotypes.

**Figure 7:**
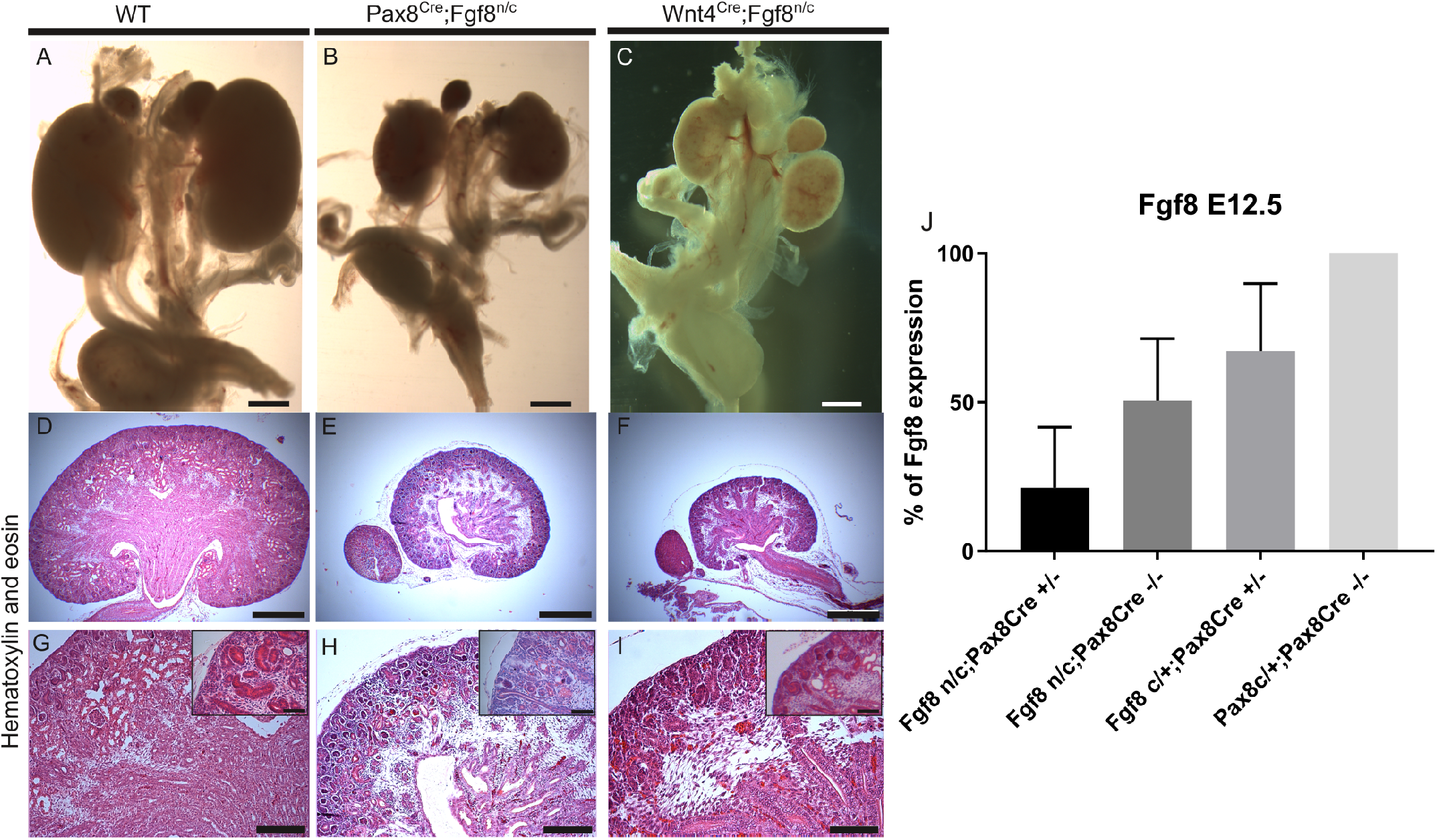
*Fgf8* deletion in E16.5 hypomorph mouse embryos. (A-C) the deletion of *Fgf8* from the metanephric mesenchyme led to smaller kidneys as compared to littermate controls. (A) Urogenital system of littermate controls, (B) Cre-mediated deletion of *Fgf8* using *Pax*8^*Cre*^ and (C) *Wnt*4^*Cre*^. (E,F,H,I) Hematoxylin-Eosin staining shows smaller kidneys lacking mature nephrons. (J) qPCR of*Fgf8* in E12.5 kidneys of *Fgf8 Pax*8^*Cre*^. The inserts are higher magnification selections. Scale bars for (A-F) are 500 μm and (G-I) are 100 μm.

### Late deletion of *Fgf8* leads to incorrect localization patterns of NPCs in the nephrogenic niche

tNPCs in the tip region of the UB are marked by *Six2* expression along with *Cited1/2* and *Eya1* (Figure 1) [2, 23, 53, 54]. *Wnt9b* that is expressed and secreted by the UB regulates the transition from tNPCs to cNPCs (Figure 1) [12]. This transition is marked by a decrease of *Cited1/2* expression [2, 14, 23]. At the same time, the expression of *Wnt4* in an aggregated subset of cNPCs that are located at the tip-trunk interface of the UB indicates the onset of nephron formation (Figure 1) [11, 18]. When we compared the expression of *Six2* in wild type kidneys to tissuespecific deletions of *Fgf8* in *Fgf* 8^*n/c*^;*Pax*8^*Cre*^ (MM and UB tip) or *Fgf* 8^*n/c*^;*Wnt*4^*Cre*^ (only MM) kidneys, we found untypical *Six*2^+^ expression patterns, indicating disorganized NPCs (Figure 8A-C). Deletion of *Fgf8* in *Fgf* 8^*n/c*^;*Pax*8^*Cre*^ kidneys did not seem to alter the expression of *Wnt9b* as compared to the littermate controls (Figure 8D-F) and as expected *Wnt9b* was not affected in *Fgf* 8^*n/c*^;*Wnt*4^*Cre*^ kidneys. On the other hand, the expression of *Wnt4* was decreased in both *Fgf* 8^*n/c*^;*Pax*8^*Cre*^ and *Fgf* 8^*n/c*^; *Wnt*4^*Cre*^ kidneys (Figure 8H,I) as compared to the littermate controls (Figure 8G). Further, in both *Fgf*8^*n/c*^; *Pax*8^*Cre*^ and *Fgf* 8^*n/c*^;*Wnt*4^*Cre*^ kidneys, condensation of NPCs expressing *Eya1* and *Cited1* around the UB was perturbed, while the expression of *Cited1* was still maintained (Figure 8K-L,N-O). Together, these data show an incorrect localization pattern of NPCs in kidneys where *Fgf8* is deleted via either *Pax*8^*Cre*^ or *Wnt*4^*Cre*^ mouse lines.

**Figure 8:**
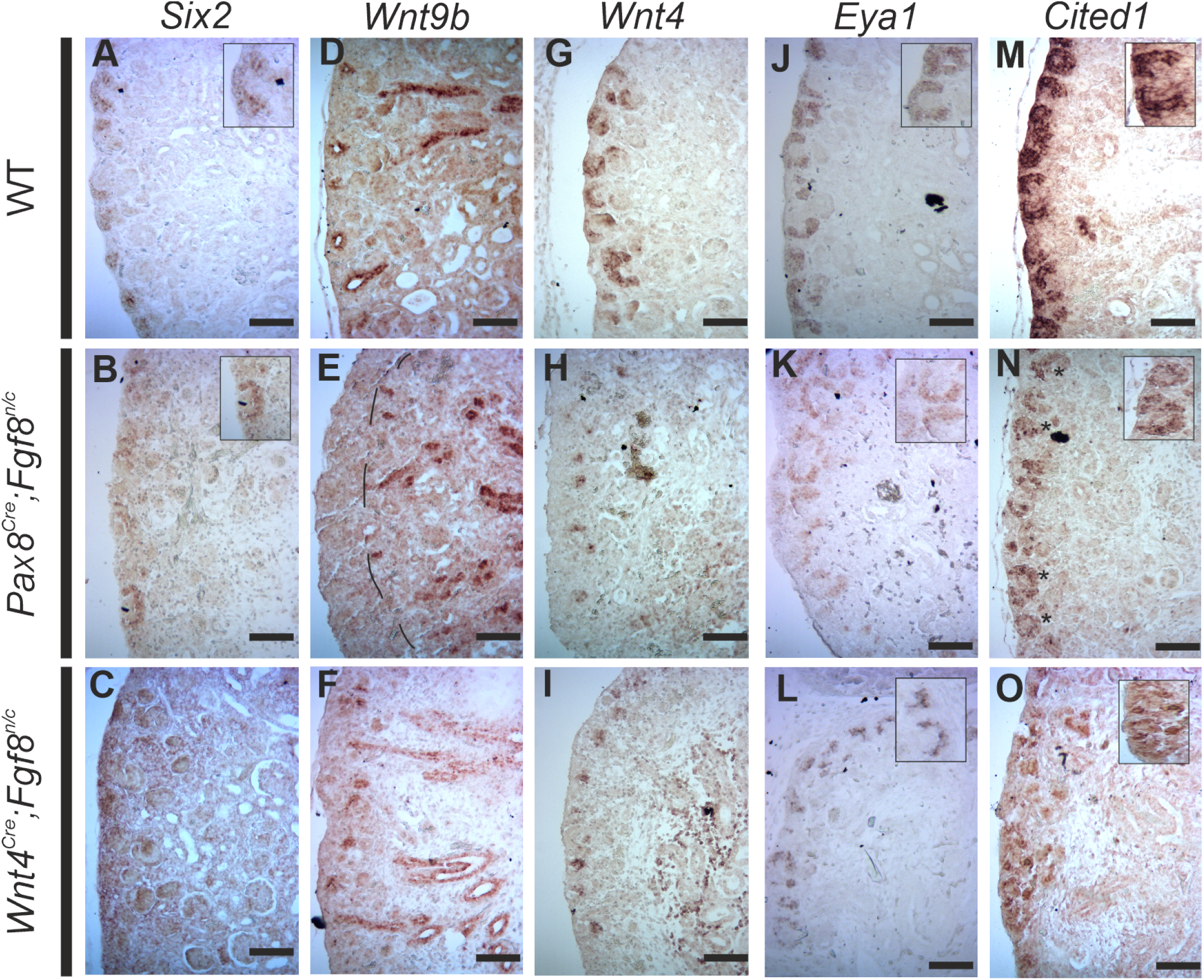
Gene expression in littermate control and hypomorph mutant kidneys.(A-C) Expression of SIX2 was observed around UB tips on the kidney cortex, but in mutant kidneys under *Pax8* or *Wnt4* Cre its expression was disorganized. (D-F) Expression of *Wnt9b* was restricted to the UB in littermate controls, *Pax*8^*Cre*^ and *Wnt*4^*Cre*^ mediated *Fgf8* deletion. (G-I) *Wnt4* expression was observed in littermate controls and mutants, but the expression was reduced in the mutants kidneys. (K-L and N-O) Expression of *Eya1* and *Cited1* was also disorganised in mutants kidneys as compared to littermate controls. Higher magnification of the staining to visualise the disorganisation of the expressing cells. Inset shows higher magnification image of the *in situ*. Scale bar represents 100 μm.

### Without the expression of *Fgf8* after kidney induction, NPCs still accumulate at the tip of the UB

To further confirm the obtained results of our *in vitro* experiments (cultured of *Fgf* 8^*n/c*^;*T^Cre^* kidneys and the effect of FGF8 onto MM cells in the bead culture experiment), we stained NPCs and UB cells with fluorescent markers on these two mouse lines, that deletes *Fgf8* later in nephrogenesis after E11.5. To analyse the complete PTAs formation, we have selected E16.5 stage where most of the tubules are already formed. Staining of *Six*2^+^ NPCs in E16.5 kidneys obtained from crossing *Fgf* 8^*n/c*^;*Pax*8^*Cre*^ revealed a thicker cap mesenchyme at the tips of UBs, suggesting differences in niche composition as compared to littermate controls (Figure 9A-B). A thicker layer of *SIX*2^+^ cells in the tip region of the UB indicates that NPCs either failed to fully condensate around the UB or that they were not primed as cNPCs (Figure 9C). Similar results were obtained when *Fgf8* was deleted using *Wnt*4^*Cre*^ (Figure 9D-E). With *Wnt*4^*Cre*^ mediated deletion of *Fgf8*, the *SIX*2^+^ population failed to condensate around the UB tip as it can be observed in *Pax*8^*Cre*^ mediated deletion of *Fgf8* (Figure 9F). These results confirm the observations from the *in situ* hybridisation experiments (Figure 8) suggesting that the NPCs, the SIX2^+^ cells, accumulate around the UB tip and that NPC induction is interrupted failing the PTAs formation.

**Figure 9:**
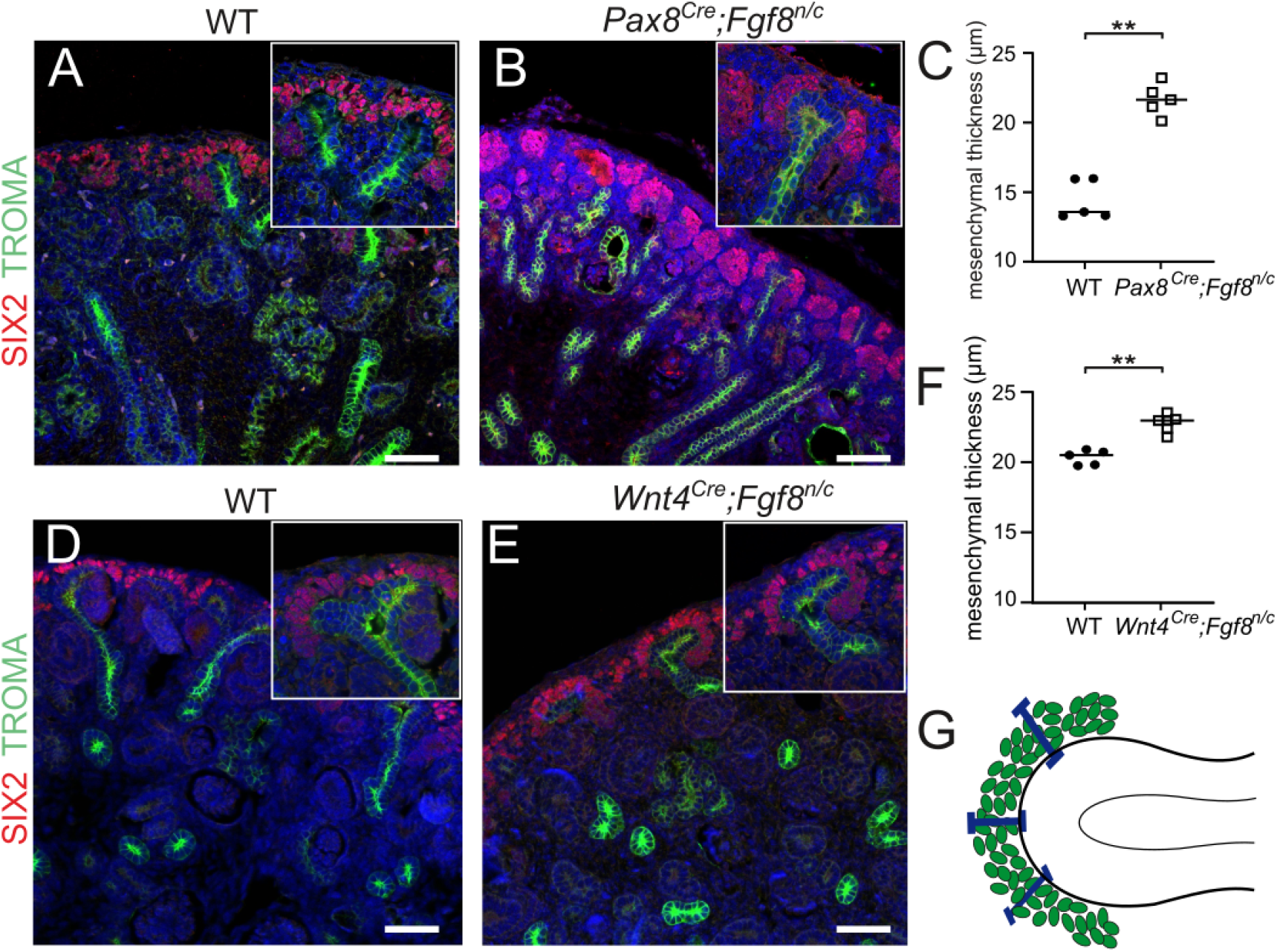
Accumulation of NPCs in the tip region in hypomorph kidneys. (A-C) When *Fgf8* was deleted using *Pax*8^*Cre*^ (B) or *Wnt*4^*Cre*^ (E), NPCs did not form a condensed cap mesenchyme, but they rather accumulated at the tip region of the UB as compared to the littermate controls (A,D), respectively. The inserts are higher magnification selections. (C) Distance of the last layer of the NPCs from the UB tip of E16.5 *Fgf* 8^*n/c*^; *Pax*8^*Cre*^ kidneys. (F) Distance of the last layer of the NPCs from UB tip of E16.5 *Fgf* 8^*n/c*^;*Wnt*4^*Cre*^ kidneys. (G) Cartoon depicting how the measurements were done for calculating the distance of NPCs around the UB tip. Kidneys were stained with SIX2 (NPCs, red), Troma-1 (UB, green) and DNA was counter stained with Hoechst33342 (Blue). Statistics were performed in the kidneys (n=6) per sample, *p* values were calculated using *t-test*, **p < 0.01. Scale bar represents 100 μm.

## Discussion

Intercellular signaling between NPCs and UB cells is key for mammalian kidney development, and it is known to be tightly controlled. Failure of the expression of key genes such as *Six2*, *Fgf8*, *Wnt4*, *Wnt9b* and others leads to developmental defects or even to embryonic lethality [2, 3, 18, 25, 26]. FGF8 is also known to be involved in the activation of the kidney-specific genes *Wnt4* and *Lim1(Lhx1)* [25, 27].

In this work, we have used several cre-based mouse lines to establish that *Fgf8* expression is located in the MM and imparts its function on the nephron progenitor cells. Deletion of *Fgf8* from the MM resulted in embryonic kidneys that lacked mature nephrons, which led to smaller hypomorphic kidneys and postnatal death. As *Fgf8* -deficient kidneys lack *Wnt4* expression, *Fgf8* is required for *Wnt4* expression [25, 26]. While it is known that WNT4 initiate MET [18], in accordance with previously published results, we also observed that independently of WNT4, FGF8 is required for the condensation of NPCs [25]. We found that in kidneys where FGF8 signaling was blocked using an anti-FGF8 antibody, NPCs did not condensate to form pretubular aggregates. But upon removal of the antagonising agent, PTA formation was recovered. This indicates that even though FGF8 is upstream of WNT4, it regulates NPC condensation, which is itself required for PTA formation before WNT4-induced MET of PTAs.

FGFs are also involved in the maintenance of NPCs during early nephrogenesis [31]. Though previously only a modest effect of FGF8 on cap marker transcription has been observed in 2D cultures [31], our results with 3D cultures show that without FGF8, *Six2* expression is lost, but when adding ectopic FGF8 and in the presence of the UB, the expression of *Six2* is maintained as compared to the vehicular control. Similarly, the expression of other tNPC markers such as *Cited1/2* and *Eya1* is also maintained in the presence of FGF8. We explain the discrepancy with the previous observations through improved culture conditions, as it has been shown that culturing NPCs in a 3D micro-environment leads to an improvement in nephrogenic potential [55]. In conclusion, FGF8 seems to be required for the expression of tNPC markers and thus tNPC maintenance, though it might not determine NPC fate, as cNPCs that form PTAs also express FGF8.

To understand how FGF8-induced NPC motility could lead to NPC condensation, we employed a computational model that simulates the combined effect of FGF8-induced motility, SHH-based repression of *Fgf8* and weak chemoattraction by WNT11. Our simulations showed that, on the one hand, graded motility perpendicular to the UB surface due to the repression of *Fgf8* by SHH and, on the other hand, the weak attraction of NPCs by UB-secreted WNT11 together established a trap-like region near the UB that captured passing NPCs. Importantly, with FGF8-induced NPC motility significantly more NPCs were trapped than with low motility and weak attraction. We, therefore, propose that FGF8-induced chemokinesis of NPCs, together with signals from the UB, represent a robust mechanism of NPC condensation in the nephrogenic niche. Our proposed mechanism may be involved both in the initial formation of the cap mesenchyme and the formation of pretubular aggregates at the tip-trunk interface of the UB. A better understanding of both processes requires an investigation of the spatiotemporal gradients of the involved factors during early nephrogenesis.

In conclusion, our results indicate that FGF8 is a chemokinetic agent that is expressed in the metanephric mesenchyme and is required for the condensation of NPCs while being involved in tNPC maintenance. Further work is required to reveal how FGF8 along with its receptors and inhibiting factors orchestrates NPC condensation, its involvement in PTA formation, and its influence on cell fates to form the various structures of the nephron.

## Methods

### Mouse strains and tissue collection

In this work, the mouse experiments were conducted in accordance with the Finnish and EU legislation. The Finnish National Animal Experiment Board approved all animal experiments and experiments were conducted under internal licenses issued by the Laboratory Animal Centre of the University of Oulu, Finland. To delete *Fgf8* from mouse kidneys we crossed *Fgf* 8^Δ2,3*/*+^ males with *Pax*8^*Cre/*+^ female. The progeny was genotyped and females with *Pax*8^*Cre/*+^;*Fgf* 8^*n/*+^ genotype were crossed with *Fgf*8^*Floxed/Floxed*^ male. The progeny were genotyped and embryos with genotype *Fgf* 8^*n/c*^;*Pax*8^*Cre/*+^ were selected for the experiments. Similar strategy was utilized for *Wnt*4^*eGF PCre*^/+ and *HoxB*7^*Cre/*+^. To delete *Fgf8* using *T^Cre^*, a similar strategy was employed as in Perantoni *et al.* [25]. Briefly, females with genotype *Fgf* 8^*Floxed/Floxed*^ were crossed with males with genotype *T^Cre/Cre^*;*Fgf* 8^Δ2,3*/*+^ and progeny with genotype of *T^Cre/^*^+^;*Fgf* 8^*n/c*^ was used for this study. Expression of *Cre* and deletion of *Fgf8* were assessed by genomic PCR as described in respective articles. Timed matings were checked at noon for vaginal plug and upon identification was considered to be E0.5. To obtain kidney samples, pregnant females were euthanized with *CO*2 followed by cervical dislocation, as per the institutional guidelines. Embryos were collected in sterile PBS and kidneys were dissected in 1X PBS with calcium and magnesium. Dissected kidneys were further treated depending on the experiment.

### Histology and immunofluorescence

#### Hematoxylin-Eosin staining

For light microscopy, paraffin-embedded kidney sections were stained with hematoxylin-eosin, following standard procedures according to Veikkolainen *et al.* [60].

#### Immunostaining

Embryonic kidneys were collected and dissected at E11.5, E12.5, E16.5 and P0 from genotyped mutant and littermate controls. Samples were fixed in 4% paraformaldehyde (PFA) overnight at 4°C and dehydrated in serial dilutions with 25% ethanol *→* 50% ethanol *→* 75% ethanol *→* 100% ethanol in water with an incubating time in each step for 45mins or until the sample sinks to the bottom of the test tube. At this point, the samples were transferred in a clearing solution (xylene), twice, for 60mins each. Samples were moved to melted paraffin, thrice, for 60mins each, and then these samples were carefully embedded in paraffin blocks and stored at 4°C. Embedded samples were sectioned (14 μm) with Lecia microtome, and slides were prepared with up to 4 samples on each slide. To perform staining, sections were selected, and slides were prepared by heating at 55°C to melt paraffin. To deparaffinise, slides were incubated in xylene solution, twice, 5mins each. To remove xylene, slides were incubated in the 100% ethanol, twice, 5mins, and then hydrated in 95% ethanol *→* 75% ethanol *→* 50% ethanol in water for 5mins each. Antigen retrieval was performed 1mM EDTA-NaOH solution (pH 8.0) or 10mM Na-citrate-citric acid solution (pH 6.0) in pressure cooker for 10mins [61]. After antigen retrieval samples were incubated, for 60 mins, in blocking solution (1X PBS + 0.01% Triton-X100 + Serum (Serum was selected based on antibody reactivity)). Primary antibody incubation duration and temperature was. Images were accquired on Olympus BX51WI upright mircroscope with Hamamatsu ORCA-ER digital camera.

#### Nephrospheres

Sphere-Forming Assay was performed as per Ihermann-Hella *et al.* with minor modifications [32]. MM from E11.5 kidneys from CD-1 mice were dissociated with Collagenase Type 3 (Worthington Biochemical Corporation; Cat #LS004180) and DNase I (New England Biolabs; Cat #M0303S) in physiological buffer at 37°C. To obtain a single-cell suspension, after stopping the activity of the enzymes with complete media, the cell suspension was strained through 0.45 μm cell strainer. The total obtained cell suspension was divided into four equal parts, from which each part was mixed with two parts of Matrigel (Corning; Cat #354277) and were allowed to attach for 10mins at 37°C, 5% *CO*2 and 95% humidity. Media containing Src-kinase inhibitor (10*μM* PP2; Tocris Bioscience; Cat #1407) and/or FGF8b (100ng/ml; R&D Systems; Cat #423-F8) 24hrs.

### Real time qPCR

To remove nephrospheres from the embedded matrigel, plates were chilled on ice for one hour on a shaking platform. Liquefied matrigel solution was collected and centrifuged at 10621 r.c.f at 4°*C* for 10 min. Pellet was washed twice with DEPC treated 1XPBS at 2655 r.c.f at 4°*C* for 5 min. Pellet was flash-frozen in liquid nitrogen until required. RNA extraction was performed using RNeasy mini (Qiagen; Cat# 74104) and cDNA synthesis was performed using First Strand cDNA synthesis kit (Thermo Scientific; Cat# K1612), where 100ng of RNA was used as a template. cDNA was diluted 1:1 with PCR grade water and for the qPCR reaction 2*μL* cDNA, 1.2*μL* each of forward and reverse primer along with 0.6*μL* of PCR water and 5*μL* Brilliant Sybr Green III qPCR master mix (Agilent Technologies; Cat# 600882). qPCR was carried out at CFX96 Touch System (Bio-Rad) with a program 95°*C* 10mins, [95°*C* for 20sec, 60°*C* for 20sec, 72°*C* for 20 sec] for 40 cycles followed by melt curve. qPCR was performed with three biological replicates in three technical replicates.

#### Western Blotting and Antibody validation

For the validation of anti-FGF8b antibody, a Flag-tagged FGF8b clone was obtained from Genscript CloneID: OMu22892D (NCBI Nucleotide: NM 001166361.1) and was overexpressed in CHOK1 cells (ATCC: #CCL-61). The media was collected from FBS starved CHO-K1 cells expressing FGF8b and precipitated using the TCA method as described in Fic *et al* [62] and western blot was performed with controls (commercially bought rmFGF8b, cell lysate of validated protein-expressing Flag tag and CHO-K1 cell lysate).

For *GSK*3*β* quantification, nephrosphere culture was setup (Material and Mehtods). The cells were extracted as previously mentioned (Material and Method). Cells where lysed using 1X RIPA cell lysis solution (Cell Signaling; Cat#9806) supplemented with cOmplete™, Mini, EDTA-free Protease Inhibitor Cocktail (Roche; #04693159001). Protein quantity estimation was performed using Pierce™ BCA Protein Assay Kit (Pierce; Cat#23225). 50mg of total protein was loaded on to in-house prepared 12.5% SDS-PAGE separating gel and 6% stalking gel and separation was performed for 90mins at 110V at room temperature. Transfer was carried out on to NCP Porablot Membrane (Macherey-Nagel; Cat#12807411) for 90mins at 90V at 4°*C*. The membrane was blocked with 5% BSA solution and all the primary antibodies were incubated for overnight at 4°*C*, while secondary antibodies were incubated for 60mins at room temperature. The detection was performed using *LumiGLO* Reagent (Cell Signalling; Cat#7003S) on Fujifilm LAS-3000 Imager. For sequential protein detection the antibodies were striped away, by incubating the membranes with 0.2M NaOH solution for 15mins at room temperature and re-blocking with BSA. The protein quantitation was performed using ImageQuant TL8.1 (Cytiva LifeSciences).

### *In Situ* hybridization

The non-radioactive section in situ hybridization technique was performed as described Junttila *et al.* [63]. The used cDNA probes were *Wnt4*, *Wnt9b*, *Six2*, *Eya1* and *Cited1* were obtained as gifts from Prof. Thomas Carroll (University of Texas Southwestern, USA).

### Flow Cytometry

Dissected E11.5 kidneys were cultured in a Trowell culture system. After 3 days of culture with or without rmFGF8b, kidneys were dissociated into a single-cell suspension. Cells were fixed with 4% PFA and permeabilized with BD Cytofix/Cytoperm (BD Biosciences; Cat:# 554714) as per the manufacturer’s instructions. Samples were stained with primary anti-SIX2 antibody for 30 mins on ice, washed, and then stained with secondary Goat anti-rabbit AlexaFluor 488 for 30 mins on ice and washed thoroughly. Samples were scored on FACSCalibur (BD Biosciences). Three biological repeats were carried out for each condition while maintaining the protocol and template for sample acquisition.

### Cap mesenchymale quantifications

Sections of E16.5 *Pax*8^*Cre*^; *Fgf*8^*n/c*^, and E11.5 *T^Cre^;Fgf* 8^*n/c*^ kidneys were cultured for 3 days in Trowell culture, fixed, and stained with anti-SIX2, and anti-TROMA-1 and counterstained with Hoechst 33342 (Thermo Scientific; Cat# H3570). Imaging was done with Zeiss LSM 780 confocal microscope and samples were analyzed on Zen Blue (2012 edition; Zeiss). The distance between the pair of *SIX*2^+^ cells closest and most distant to the UB was measured repeatedly along the UB at intervals of one cell length to determine the thickness of the cap mesenchyme. Additionally, the dispersion of NPCs was quantified as the Euclidean distance between NPCs and UB surfaces. Using Bitplane Imaris 9.6.0 and the Imaris modules Measurement Pro and Vantage, NPC positions were quantified using fluorescence intensity-based spot detection and UB surfaces were segmented using fluorescence intensity-based 3D segmentation. To determine the proportions of attached and free/unattached *SIX*2^+^ cells in both controls and mutant kidneys, cells were classified as attached when their centroid positions were closer to the UB than twice the median (17.4 μm) of all centroid positions of the *SIX*2^+^ cells in the control kidneys.

### Bead assays with NPCs

A sphere forming assay was modified to induce cell motility in response to FGF8. Metanephric mesenchyme cells were dissociated into a single-cell suspension and an equal amount of cells were divided and mixed with Matrigel. BSA soaked or FGF8 (100 μg/ μl) soaked agarose blue beads (Affi-Gel Blue Gel, Bio-Rad Cat#1537302) were carefully placed in individual wells of the 4 Chamber 35mm glass-bottom dish (Cellvis, Cat #D35C4-20-0-N). The Matrigel cell mixture was carefully applied to the surface of the beads and placed in a pre-equilibrated microscopic chamber maintained at 37^*◦*^C and 5% *CO*2. Time-lapse was performed with Leica SP8 falcon 20X water immersion lens for 24hrs. Cells were tracked using the Imaris (v9.6; Bit-Plane, South Windsor, CT, USA) cell tracking functionality. Cell tracks from different samples (n=5 each) were pooled and analyzed using the R package *CelltrackR* [64].

### Modelling

Our model utilizes a cell-based Lattice-Boltzmann Immersed-Boundary simulation framework for morphogenetic problems [66].

#### Setup

We simulated mesenchymal condensation within a 150 *μm*^2^ section of the nephrogenic niche containing randomly dispersed NPCs and a flat patch of ureteric epithelium for an interval of 12 hours, corresponding to embryonic days E10.5 to E11 (Figure 5D, (Supplementary Movie 3). The model represents the dynamics of FGF8, SHH and WNT11 concentrations by the following differential equations:

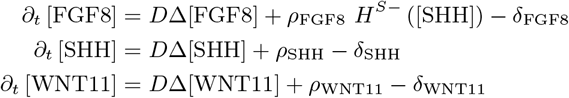

*∂_t_* is the time derivative, *D* is the diffusion coefficient, Δ denotes the Laplacian operator, *ρ, δ* are the production and degradation rates and *H^S−^* is the shifted Hill function. Parameter values are shown in Table 4.

**Table 1:** Mouse lines used in this work.

| Reagents type (Species) or resource | Designation | Source or reference | Identifiers |
| --- | --- | --- | --- |
| Genetic reagent (Mus musculus) | *Pax8<sup>Cre/+</sup>* | [50] | MGI:J:80208 |
| Genetic reagent (Mus musculus) | *Wnt4<sup>eGFPCre/+</sup>* | [52] | RRID:IMSR_EM:10011 |
| Genetic reagent (Mus musculus) | *Hoxb7<sup>Cre/+</sup>* | [49] | MGI:J:79481 |
| Genetic reagent (Mus musculus) | *Fgf8<sup>Floxed/Floxed</sup>* | [56] | MG:2150347 |
| Genetic reagent (Mus musculus) | *Fgf8<sup>Δ2,3/+</sup>* | [56] | MGI:2150346 |
| Genetic reagent (Mus musculus) | *T<sup>Cre/+</sup>* | [25] | MGI:J:101175 |
| Genetic reagent (Mus musculus) | *Rosa26<sup>mT/mG</sup>* | [57] | MGI:J:124702 |
| Genetic reagent (Mus musculus) | *Six2 – TGC<sup>tg</sup>* | [58] | MGI:J:148455 |
| Genetic reagent (Mus musculus) | *Rosa26 – LacZ<sup>loxP</sup>* | [59] | MGI:J:64292 |

**Table 2:** Antibodies, growth factors, agonists and antagonists.

| Product | Final Concentration | Company; Catalogue number; Identifiers |
| --- | --- | --- |
| Anti-SIX2 | 1:200 | Proteintech Group; Cat# 11562-1-AP;<br>RRID:AB_2189084 |
| Anti-FGF8b | 1:200 | R&D Systems; Cat# AF-423-NA; RRID:AB_2262650 |
| Anti-Flag M2 | 1:300 | Sigma-Aldrich; Cat# F1804; RRID:AB_262044 |
| Anti-TROMA1 | 1:200 | DSHB; Cat# Troma1; RRID:AB_531826 |
| Anti-*GSK3β* | 1:1000 | Cell Signalling; Cat#9315S; RRID:AB_490890 |
| Anti-phospho-*GSK3β* | 1:1000 | Cell Signalling; Cat#9336S; RRID:AB_331405 |
| Anti-β – *ACTIN* | 1:1000 | Cell Signalling; Cat#4970S; RRID:AB_2223172 |
| Goat anti-Rabbit | 1:1000 | ThermoFisher Scientific; Cat# A-11008; |
| AlexaFlour488 |  | RRID:AB_143165 |
| Goat anti-Rat | 1:1000 | ThermoFisher Scientific; Cat# A-11081; |
| AlexaFlour546 |  | RRID:AB_2534125 |
| Goat anti-Rabbit | 1:1000 | AgilentDako; Cat# P0448; |
| HRP |  | RRID:AB_2617138 |
| rmFGF8b | 100ng/μL | R&D Systems; Cat# 423-NA |
| PP2 | 10 μM | Tocris Bioscience; Cat# 1407; PubChem ID: 4878 |
| Bio | 10 μM | Tocris Bioscience; Cat# 3194; PubChem ID: 5287844 |
| Matrigel (BME) |  | Corning; Cat# 354277 |

**Table 3:**
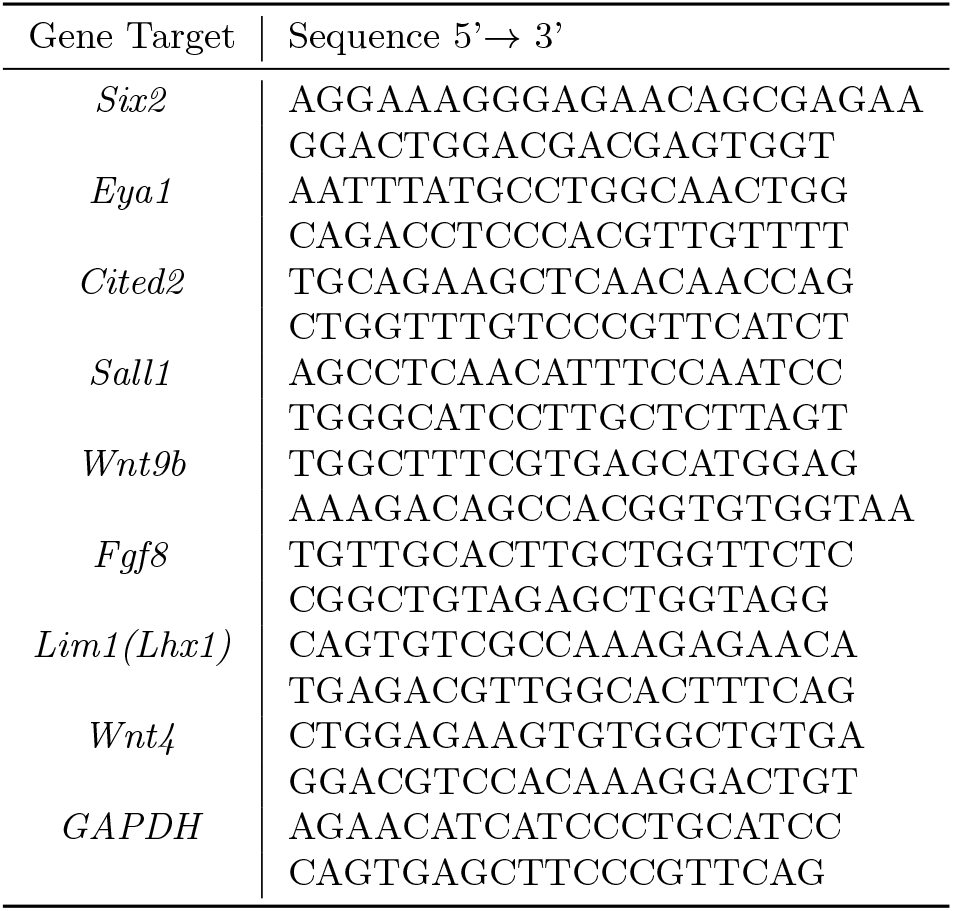
qPCR primers.

**Table 4:**
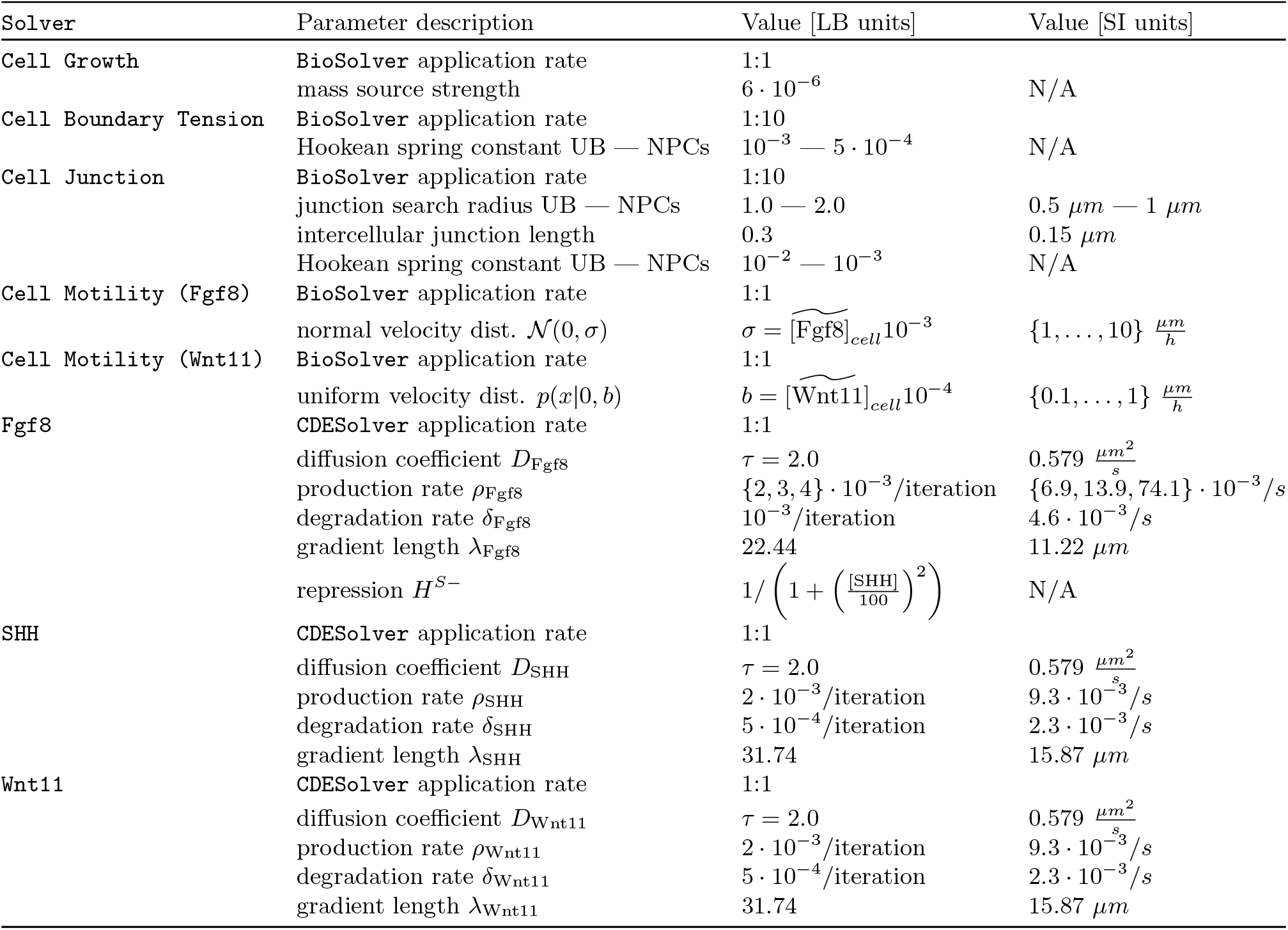
Parameter values for the simulation of mesenchymal condensation in the nephrogenic niche. 1 length unit of the simulation box corresponds to half a micron, i.e. 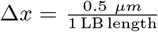 and 2 *·* 10^5^ iterations correspond to 12 hours of developmental time, i.e. 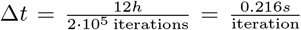. The diffusion coefficient *D* of the morphogens is related to the LB relaxation time *τ* via 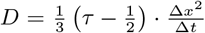 [65]. The gradient length 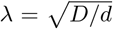 of an exponential gradient *c*(*x*) = *c*_0_*e^−x/λ^* is the distance at which the concentration has decreased to 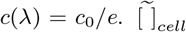 is the median intracellular concentration. Mass source strength as well as production and degradation rates refer to the PhysicalNodes which represent the discrete grid on which the fluid and the morphogens live [65]. The area of an NPC or an epithelial cell comprises a few hundred PhysicalNodes; the simulation box comprises exactly 300 *·* 300 PhysicalNodes. Morphogen dynamics are described above, under *Setup*.

#### Measurements

At the end of each simulation, the distance of all NPC centroids to the centroid of the ureteric epithelium was measured. All data were pooled and visualized as a boxplot (Figure 5C) using R [67]. The significance level was determined based on a Wilcoxon signed-rank test and added to the boxplot.

#### Cells

Cells are represented by highly resolved 2D polygonal geometries with a cortical tension established by elastic forces between GeometryNodes [66]. Similarly, cell adhesion is realized by elastic forces between neighbouring cells. GeometryNodes are added or removed when the distance between GeometryNodes, i.e. cell size, changes. Similarly, adhesions are created or removed based on a distance threshold between intercellular GeometryNodes. The cells are immersed in a Newtonian fluid and no-flux boundary conditions are imposed on the domain boundaries.

#### Morphogens

Morphogens are produced and degraded within cells, can freely diffuse within the entire domain and through cell boundaries and are advected by motile cells. Parameter values are shown in Table 4.

#### Cell motility

Random motility is established by applying 2D velocities to each GeometryNode, where velocities are picked from a normal distribution whose standard deviation is proportional to the median local FGF8 concentration (Figure 5B). Similarly, a weak attractive force is established by picking positive 1D velocities (directed towards the UB) from a uniform distribution where the upper bound is proportional to the median local WNT11 concentration.

## Acknowledgments

We thank Ms Paula Haipus, Ms Johanna Kekolahti-Liias, and Ms Hannele Härkman for excellent technical assistance, and Prof Maxime Bouchard for providing *Pax*8^*Cre*^, *Fgf* 8^*Floxed/Floxed*^, and *Fgf* 8^Δ2,3*/*+^ and Prof Andy McMahon for *Hoxb*7^*Cre*^ mouse line. We also would like to thank Tiina Jokela for collecting the mouse samples and establishing the protocol for genotyping. We would like to thank Dr. Alan O Perantoni and Dr. Mark B Lewandoski for providing us with *T^Cre^*;*Fgf* 8^*n/c*^ mouse embryonic kidneys. Dr Veli-Pekka Ronkainen is acknowledged for help in setting up the time-lapse imaging, and the Vainio lab for helpful discussions. Marco Meer thanks Harold Gomez for discussions on image analysis and Lisa Conrad for comments on the manuscript. Confocal imaging was conducted at the Light Microscopy Unit of Biocenter Oulu, University of Oulu. This work was funded for Prof Seppo Vainio by the Academy of Finland (206038, 121647, 250900, 260056), Centre of Excellence Grant 2012–2017 of the Academy of Finland (251314), and Tekes BioRealHealth (24302443); for Dr Florence Naillat, by the Academy of Finland post-doctoral Fellowship (243014583), the Foundations: Post Doc Pool (Svenska Kulturfonden) and the Finnish Cultural Foundation (Pekka ja Jukka-Pekka Lylykarin rahasto); for Marco Meer and Prof Dagmar Iber by the SNF Sinergia grant CRSII5 170930.

## Author Contributions

AS, SV & FN conceived the study, AS, AD & FN carried out the experiments and acquired the data, AS, MM & FN analyzed the data and wrote the manuscript, MM developed the computational model with help by DI and carried out the simulations, and AS & MM produced the figures. All authors have read the manuscript and approved it for publication.

## Supplementary Material

Supplementary movies and source code are available as a git repository^*^.

## Supplementary Figures

**Figure S1:**
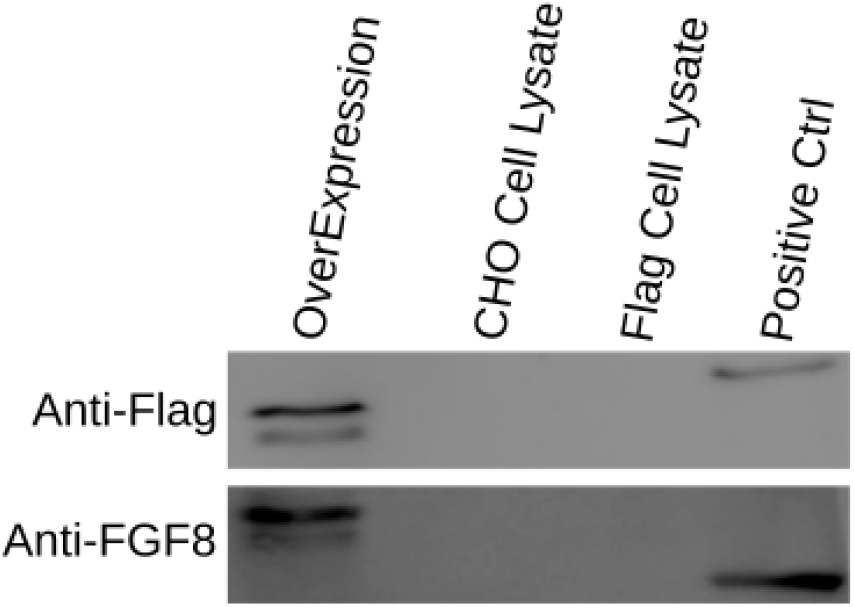
The specificity of the FGF8 antibody was measured using flag-tagged FGF8 were expressed in CHO cells. Recombinant mouse FGF8b was used as a positive control.

**Figure S2:**
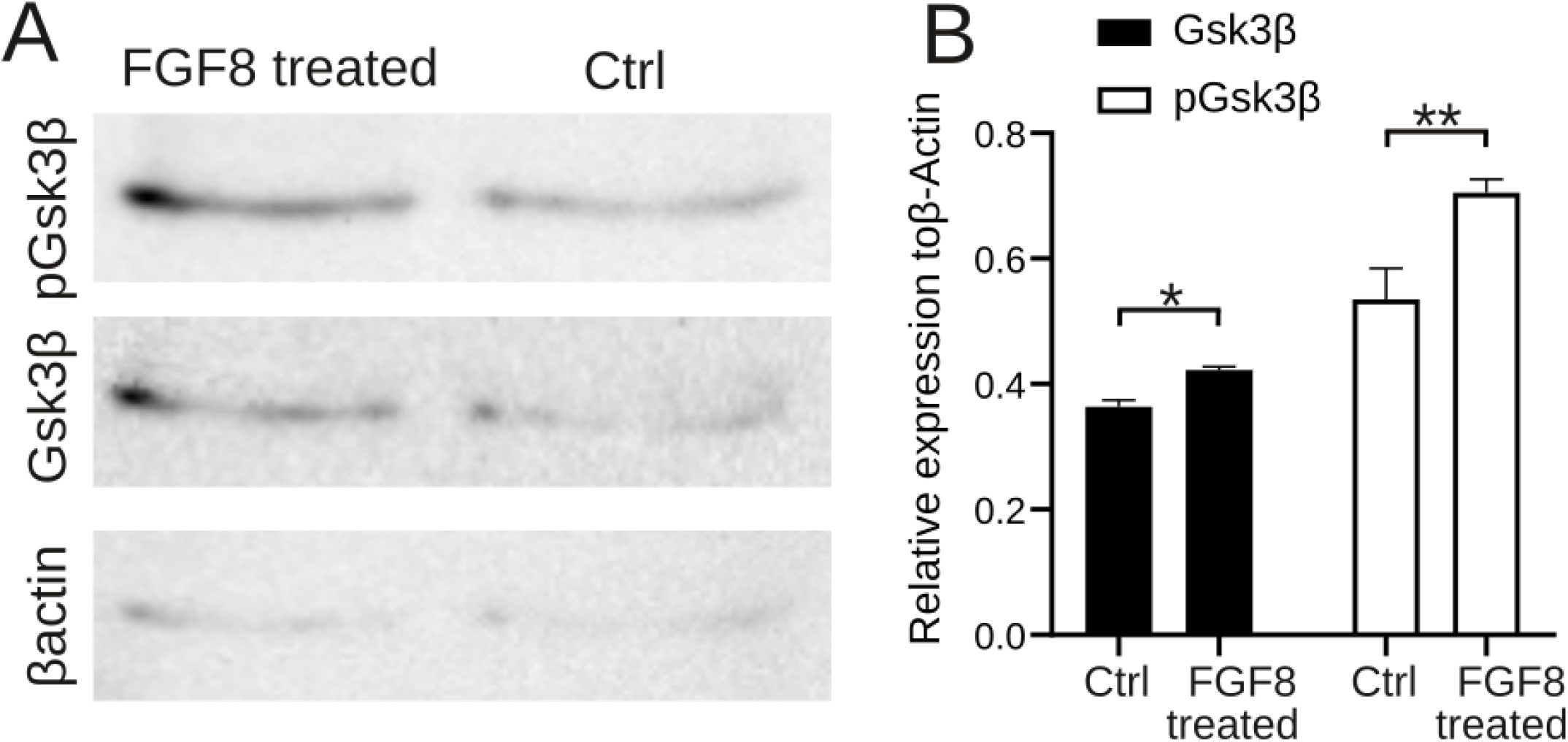
Expression of *GSK*3*β* in response to Fgf8b.A) Western blot analysis of phospho-*GSK*3*β* production in response to ectopic FGF8b in nephrosphere culture. B) Relative expression of *GSK*3*β* and phospho-*GSK*3*β*. Statistics were performed (n=3) using Two-way ANOVA. **p < 0.023 and *p < 0.028.

**Figure S3:**
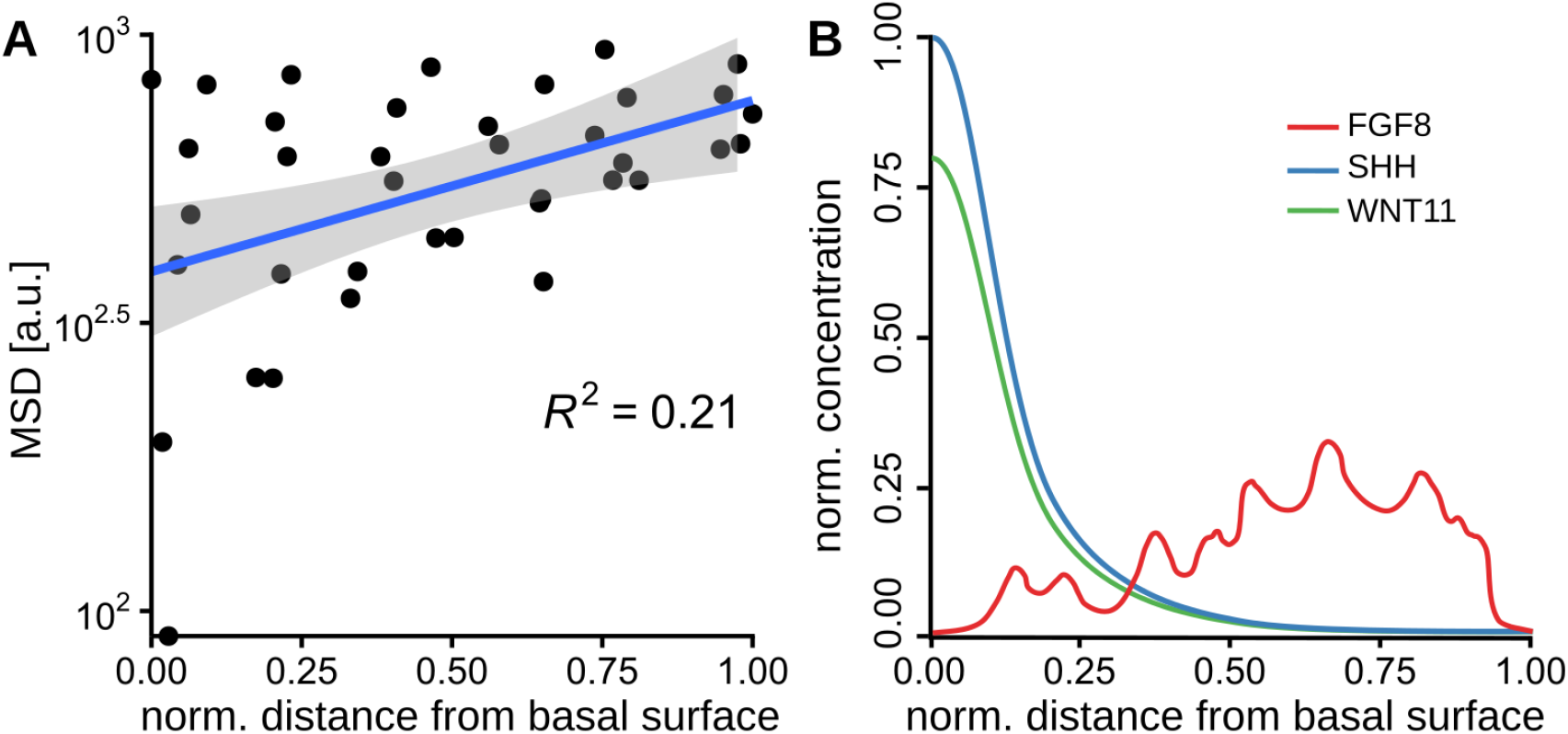
Simulated motility and concentration gradients. (A) The observed motility gradient in the simulations quantified by the mean squared displacement (MSD) of each NPC from its initial position with respect to the distance from the basal surface of the ureteric epithelium. Each dot represents the average over n=20 simulations of the squared displacement of each NPC from the same initial position. The blue solid line and shadowed region represent a linear regression with confidence interval (95% confidence level). The low *R*^2^ indicates the variability of NPC displacements, reflecting the randomness of NPC motility. (B) Example of concentration profiles in the simulations. Curves represent concentration profiles along a line perpendicular to the basal surface of the ureteric epithelium. Concentrations were normalized to the maximum SHH concentration.

**Figure S4:**
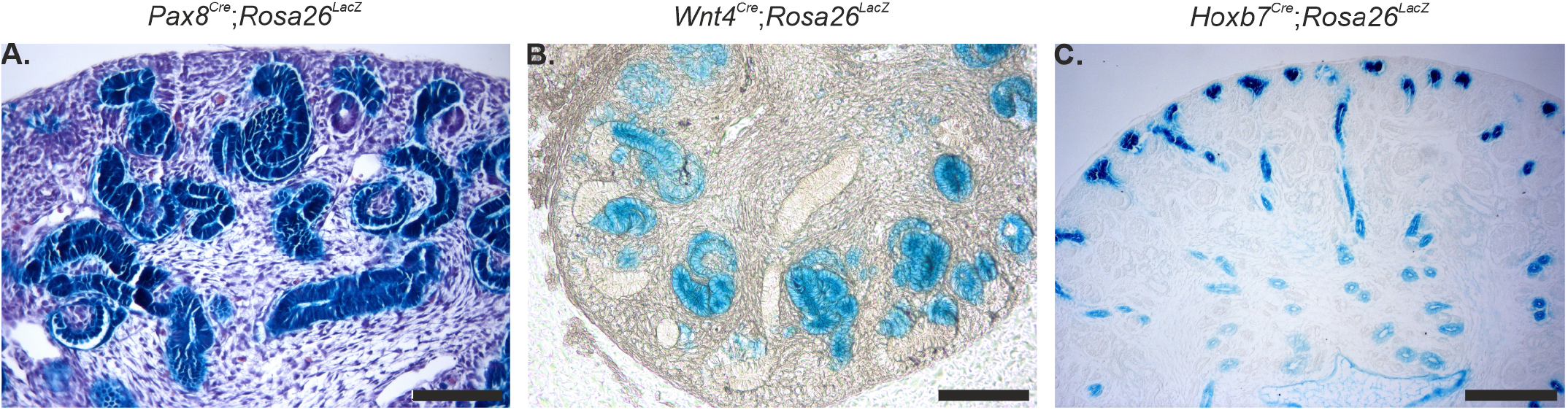
X-gal staining after *β Galactosidase* expression due to Cre recombination. A) E14.5 *Pax*8^*Cre*^, B) *Wnt*4^*Cre*^ and E16.5 *Hoxb*7^*Cre*^ mouse kidneys. A) Expression of PAX8 is limited to the UB tip and MM. B) WNT4 expression is observed in PTAs. C) HOXb7 expression is limited UB and no expression was observed in the MM. Scale bar represents 100 μm.

**Figure S5:**
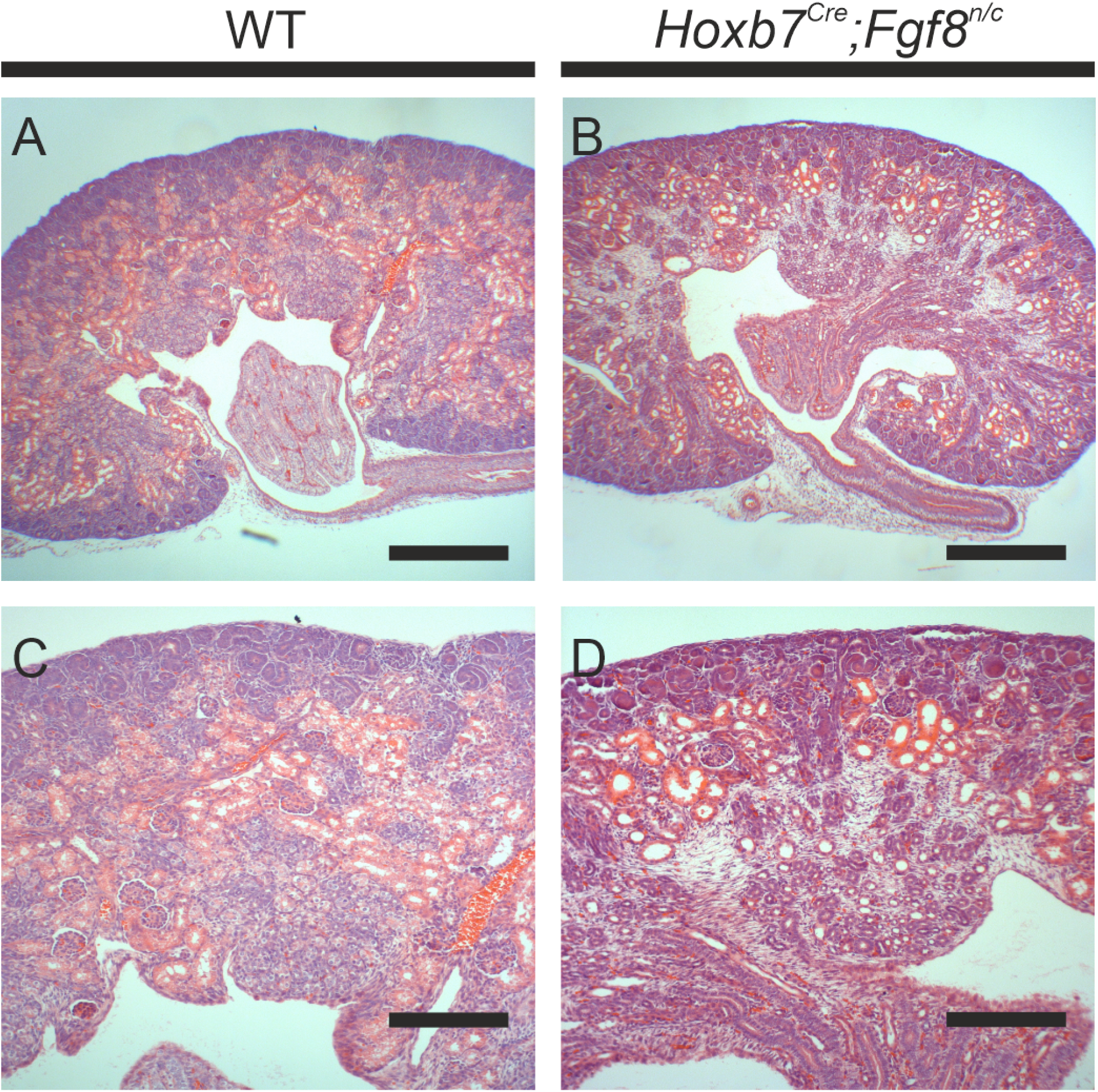
Hematoxylin-Eosin staining of (A-C) littermate control (WT) and (B-D) *Fgf* 8^*n/c*^;*Hoxb*7^*Cre*^ shows no phenotype. Scale bar represents 100 μm.

## Supplementary Movies

**Supplementary Movie 1:**

https://git.bsse.ethz.ch/iber/Publications/2022_Meer_NPC_Condensation/-/blob/main/Video1_spherification_montage.mp4. 24-hour time lapse imaging of *in vitro* experiments where NPCs were cultured in a 3D matrix. A) NPCs cultured without any additive (control), B) NPCs cultured in FGF2, C) NPCs cultured in FGF8 and D) NPCs cultured in anti-FGF8b antibody. Scale bar represents 50 μm.

**Supplementary Movie 2:**

https://git.bsse.ethz.ch/iber/Publications/2022_Meer_NPC_Condensation/-/blob/main/Video2_cell_motility_quant.mp4. Time lapse imaging of *in vitro* experiments with control beads (A-E) vs. FGF8 soaked beads (F-J), 5 samples each. Dragon tails are shown. Scale bars: 50 μm.

**Supplementary Movie 3:**

https://git.bsse.ethz.ch/iber/Publications/2022_Meer_NPC_Condensation/-/blob/main/Video3_mesenchymal_condensation_simulations.mp4. Simulations of increasing relative FGF8 concentrations: Scarce, intermediate (2x increased) and high (3x increased).

## Footnotes

∗ https://git.bsse.ethz.ch/iber/Publications/2022_Meer_NPC_Condensation

